# Preclinical Development of a Romidepsin Nanoparticle Demonstrates Superior Tolerability and Efficacy in Models of Human T-Cell Lymphoma and Large Granular Lymphocyte Leukemia

**DOI:** 10.1101/2024.07.18.603379

**Authors:** Ipsita Pal, Anuradha Illendula, Andrea Joyner, John Sanil Manavalan, Tess M. Deddens, Ariana Sabzevari, Deepthi P. Damera, Samir Zuberi, Enrica Marchi, Todd E. Fox, Marya E. Dunlap-Brown, Kallesh D. Jayappa, Jeffrey W. Craig, Thomas P. Loughran, David J. Feith, Owen A. O’Connor

## Abstract

Histone deacetylase (HDAC) inhibitors are a widely recognized and valued treatment option for patients with relapsed or refractory peripheral T cell lymphomas (PTCL). Romidepsin is a relatively selective Class I HDAC inhibitor originally approved for patients with relapsed or refractory (R/R) cutaneous T cell lymphoma (CTCL) and subsequently R/R PTCL. Unfortunately, the FDA approval of romidepsin for R/R PTCL was withdrawn due to a negative Phase 4 post-marketing requirement (PMR), diminishing further the treatment options for patients with PTCL. Herein we describe the development of a first-in-class polymer nanoparticle of romidepsin (Nanoromidepsin) using an innovative amphiphilic di-block copolymer-based nanochemistry platform. Nanoromidepsin exhibited superior pharmacologic disposition, with improved tolerability and safety in murine models of T-cell lymphoma. Nanoromidepsin also exhibited superior anti-tumor efficacy in multiple models including *in vitro* T cell lymphoma (TCL) cell lines, *ex vivo* LGL leukemia primary patient samples, and murine TCL xenografts. Nanoromidepsin demonstrated greater accumulation in tumors and a statistically significant improvement in overall survival (OS) compared to romidepsin in murine xenograft models. These findings collectively justify the clinical development of Nanoromidepsin in patients with T-cell malignancies.

## Introduction

The histone deacetylase inhibitors (HDACi) are widely recognized as an important class of drugs for the treatment of T-cell malignancies. To date, four HDACi have been approved globally for the treatment of patients with relapsed or refractory (R/R) cutaneous (CTCL) and peripheral T-cell lymphomas (PTCL). Vorinostat, a hydroxamic acid derivative, was the first HDACi to be approved for the treatment of any disease, being approved in 2006 for the treatment of R/R CTCL (*1, 2*). Romidepsin is a bicyclic peptide originally approved for patients with R/R CTCL and PTCL in 2011 and 2014 respectively(*3–5*). Belinostat and tucidinostat (formerly known as chidamide), both hydroxamic acid derivates, were approved for R/R PTCL in 2014 (*6*), with tucidinostat being approved only by the National Medical Products Administration (NMPA, formerly the CFDA) in China (*7*). Tucidinostat has also received regulatory approval in China in combination with an aromatase inhibitor for post-menopausal women previously treated with an endocrine therapy who have hormone positive, HER2 negative recurrent or locally progressive advanced metastatic breast cancer (*8*). Panobinostat, a very potent hydroxamic acid HDACi with early promising experiences in T-cell neoplasms (*9*), was approved in 2015 in combination with bortezomib for patients with R/R multiple myeloma. Panobinostat was subsequently withdrawn in 2021 after the sponsor concluded the Phase IV commitment was not feasible (*10*). Across patients with R/R PTCL, recognizing that there are no randomized studies of one HDACi against another, these drugs produce overall response rates of approximately 25% with a progression free survival of 3-4 months with a duration of response of approximately one year (*3, 5–7*). While these drugs appear to benefit patients across the nearly 36 different subtypes of PTCL, there is some retrospective data to suggest that patients with PTCL of a T-follicular helper phenotype (PTCL-TFH) and angioimmunoblastic T-cell lymphoma (AITL) may have slightly greater benefit from an HDACi, with some studies noting the described aberrations in epigenetic genes as one possible explanation for the increased sensitivity (*11*).

While there is little dispute that the HDACi can produce meaningful cytotoxicity across a variety of cancer cell lines, clinically the drugs appear to have a unique therapeutic benefit in patients with T-cell lymphomas, though there are no compelling explanations for this disease specific activity. Histone deacetylases catalyze the deacetylation of histone and non-histone proteins. Deacetylation of histone leads to the condensation of chromatin (to heterochromatin) and transcriptional repression (*12*). Inhibition of histone deacetylases prevents deacetylation of key histones like histone 3 (H3) and histone 4 (H4), promoting open chromatin (euchromatin) and transcriptional activation. There are 11 distinct types of histone deacetylases, often categorized into class I, IIA, IIB, III and IV. Class III HDACs are not affected by any of the available HDACi, and are often referred to as sirtuins (Sirt) which are known to mediate deacetylation of proteins like p53. Class I HDACs typically comprise HDACs 1, 2, 3 and 8; Class IIA include 4, 5, 7 and 9; Class IIB include 6 and 10, and Class IV is comprised of only HDAC 11 (*13*). Many studies have sought to characterize the various HDACi as being selective for one or another of these HDAC classes or specific isoforms, though at the relatively high plasma concentrations achieved in patients, the majority of the selectivity is lost. Romidepsin exhibits nanomolar potency against class I HDACs, while the hydroxamic acids generally have a broader pattern of activity and can be more accurately defined as pan-HDAC inhibitors, inhibiting Class I, II and IV(*14*). While the Kd of any given HDACi against a particular isoform may vary, it is also clear that the profiles of genes activated or repressed by the different HDACi can vary significantly as a function of the HDACi, its concentration, its duration of exposure and the disease specific context. Efforts to ascribe inhibition of a particular HDAC to some predictive biomarker or outcome metric have been largely unsuccessful. As such, these drugs are often considered pleiotropic, and induce a broad spectrum of cellular effects. Complicating the mechanism of action further has been the recognition that HDACs can also deacetylate a host of non-histone proteins like Bcl-6, heat shock protein and others(*15*). The implications of these effects in any given disease are presently unclear.

Despite the reproducible activity of these drugs in patients with R/R PTCL, a recent Phase IV post-marketing commitment of Romidepsin-CHOP Vs. CHOP reported no difference in progression free survival (PFS) or overall survival (OS) between the arms, compelling the sponsor and U.S. FDA to withdrawal the PTCL indication for romidepsin (*16*). This, coupled with the recognition that pralatrexate is now off patent in the U.S., is leading to reduced access to important drugs used to manage patients with R/R PTCL. In general, chemotherapy regimens, especially those modeled after the paradigms deployed to treat R/R aggressive B-cell malignancies, produce excessive toxicity with minimal clinical benefit in the T-cell lymphomas. As such, the options to treat patients with R/R PTCL are dwindling, leaving physicians and patients with few to no compelling alternatives.

Nanomedicine is a field that has been rapidly growing over recent years. Nanoparticle-based drug delivery systems offer the prospect of improving pharmacokinetic profile, tissue specificity and tumor penetration, resistance to premature enzymatic degradation, increased drug retention, and reduced systemic toxicities (*17*). Liposomes have been the most common method for generating therapeutic nanoparticles for anti-cancer applications. These drugs have generally been shown to have superior tolerability and efficacy, while also exhibiting improved pharmacologic features(*18–20*). However, liposomal lipids are subject to oxidation and liposomes face challenges in encapsulating therapeutic doses of many drugs, including hydrophobic ones and especially hydrophilic pharmaceuticals. (*21*). The development of amphiphilic block co-polymer nanoparticles has markedly expanded the repertoire of drugs that can leverage the advantages of a nanoparticle mediated drug delivery (*22*). These polymers offer biocompatibility, increased stability and superior versatility in the types and combinations of drugs that can be encapsulated. We sought to synthesize a nanoparticle of romidepsin that might overcome some of the historic liabilities associated with the drug, while capitalizing on the benefits of a novel nanochemistry platform. Herein we report on the development of the first polymer nanoparticle (PNP) of romidepsin, Nanoromidepsin, demonstrating the superior properties of the molecule in contrast to the ‘naked’ version of romidepsin that has been used clinically for decades. Our approach was to design a di-block co-polymer nanoparticle with minimal excipients, in a highly controlled, scalable fabrication process using materials that are already used in clinical settings.

## Materials and Methods

### Cell Lines

The T cell lymphoma cell lines HH and H9 (both cutaneous T cell lymphomas; CTCL), SUP-T1 (T cell Lymphoblastic Lymphoma) and SUP-M2 (ALK-negative anaplastic large cell lymphoma; ALCL), were obtained from ATCC. FEPD is a gift from Dr. Salvia Jain. NKL (natural killer cell lymphoblastic leukemia/lymphoma) cell line was kindly provided by Dr. Howard Young at the National Cancer Institute. The cutaneous melanoma cell line FM3-29 was obtained from DSMZ. The Large Granular Lymphocyte (LGL) leukemia cell line TL-1 was generated in the lab of Dr. Thomas Loughran (*23*). All cells were grown at 37°C and 5% CO2 in a humidified incubator. Cell lines were authenticated by short tandem repeat DNA profiling (Genetica DNA laboratories) and tested for mycoplasma contamination routinely using the MycoAlert PLUS detection kit (Lonza #LT07–710) and Mycoplasma detection kit from Southern Biotech. Experiments were performed within 6 weeks of thawing. HH, H9, SUP-T1, FEPD and FM3-29 cells were cultured in RPMI-1640 (Corning, Glendale, AZ) with 10%-20% FBS (Thermofisher Scientific, Waltham, MA); SUP-M2 cells were cultured in RPMI-1640 with 20% FBS; TL-1 cells were cultured in RPMI-1640 with 10% FBS and supplemented with 200 U/mL IL-2 (Miltenyi Biotec cat # 130-097-743) and NKL cells were cultured in RPMI-1640 with 10% FBS and supplemented with 100 U/mL IL-2.

### Chemicals

Romidepsin was purchased from eNovations Chemicals LLC, mPEG-PDLLA and mPEG PLGA were purchased from AKiNA PolySciTech, 3,3’ -dioctadecyloxacarbocyanine perchlorate (DiO) was procured from Cayman chemical company and Poloxamer 188 was acquired from Sigma Aldrich. Acetonitrile was purchased from Fisher Scientific. Amicon ultra centrifugal filters of MWCO of 30kDa, 50kDa and 100kDa were purchased from Millipore, Sigma.

### Fabrication of Nanoromidepsin

We adopted a tandem parallel synthesis approach in order to achieve optimal physicochemical properties (>500 µg/mL romidepsin, <100 nm particle size, and <0.2 polydispersity index (PDI)) using a versatile nanoprecipitation method (WO202306463). We explored the influence of selected parameters of the nanoprecipitation method including, but not limited to, solvent to anti-solvent ratio and drug to polymer ratio to produce romidepsin loaded nanoparticles meeting the pre-determined criteria. Briefly, Nanoromidepsin was synthesized using romidepsin, diblock copolymers either mPEG-PDLLA or mPEG-PLGA, and the surfactant poloxamer 188. Constituents were dissolved in acetonitrile and added dropwise at a defined rate by a syringe pump into water while stirring. Solutions were left to evaporate for 3-4 hours to allow nanoparticle assembly. The nanoparticle colloidal solution was processed to remove un-encapsulated drug and polymer by using 100 kDa molecular weight cut-off centrifugal filters at 1460 relative centrifugal force (rcf) at 25°C for 20-60 minutes. The purified samples were collected from the filter and reconstituted into ultrapure Milli-Q water or PBS (Nanoromidepsin PDLLA H_2_O and Nanoromidepsin PDLLA PBS respectively), and analyzed using dynamic light scattering (DLS) and mass spectroscopy. For biodistribution studies, DiO, a fluorescent hydrophobic dye, was dissolved in DMSO and used as a stock solution and diluted with acetonitrile to a concentration of 1 mg/mL, then mixed with the drug and polymer solution. The drug to dye ratio was maintained at a ratio of 10:1 w/w. Co-loaded Nanoromidepsin-DiO polymer nanoparticle and ghost-DiO polymer nanoparticles were prepared using a similar method as described above.

### Encapsulation Efficiency

The encapsulation efficiency (EE) of romidespsin was determined after the romidepsin PNPs were dissolved in methanol. The concentration of encapsulated romidepsin was estimated using LC-MS/MS. The following equation was used to calculate the encapsulation efficiency: EE (%) = (Weight of the drug loaded in the polymeric nanoparticles)/ (Weight of the drug initially used in the fabrication of the nanoparticle) x 100

### Dynamic Light Scattering

Hydrodynamic size (diameter) and Polydispersity Index (PDI) of the PNP was measured in aqueous solutions using dynamic light scattering (DLS) (Malvern Instruments model ZEN 3690, Malvern, Worcestershire, WR141XZ, United Kingdom) at 25°C. This measurement includes the intensity-weighted average diameter of the particles (Z-avg), PDI, the volume-weighted average diameter over the major volume peak (Vol-Peak) and its percentage of the total population (Vol-Peak %Vol).

### Electron Microscopy

The size and morphology of Nanoromidepsin and ghost PNP were measured using a FEI Tecnai F20 (FEI, Hillsboro, OR) transmission electron microscope operating as a 120 kV cryo-Electron microscopy (cryo-EM). The cryo-EM samples were prepared using a standard vitrification method. An aliquot of ∼3 μl sample solution was applied onto a glow-discharged perforated carbon-coated grid (2/1-3C C-Flat; Protochips, Raleigh, NC, USA) where the excess solution was blotted with filter paper. The samples were then quickly plunged into a reservoir of liquid ethane at −180°C. The vitrified samples were stored in liquid nitrogen and transferred to a Gatan 626 cryogenic sample holder (Gatan, Pleasentville, CA) and then maintained in the microscope at −180°C. All images were recorded with a Gatan 4K x 4K pixel CCD camera under cryo-condition at a magnification of 9,600X or 29,000X with a pixel size of 1.12 nm or 0.37 nm, respectively, at the specimen level, and at a nominal defocus ranging from −1 to −3 μm. The unfiltered samples were recorded at 9,600X.

### Liquid Chromatography and Mass Spectrometry for Romidepsin Quantification

Romidepsin D7 is a deuterated version of romidepsin and was used as internal standard for romidepsin quantification. Romidepsin-d7 was spiked into 25 mL of plasma and the protein precipitated by the addition of 100 μl of cold 5:3 methanol: acetonitrile and centrifuged for 10 min at 4°C and 13000g. The supernatant was subsequently analyzed by liquid chromatography-mass spectrometry. Liquid chromatography was performed on an I-class Acquity (Waters, Milford, MA) with an Acquity BEH C18 2.1 x 50 mm 1.7 µm column maintained at 50°C. The mobile phases were water or methanol, each with 0.1% formic acid. The flow (0.5 mL/min) was maintained at 5% methanol for 5 minutes before a linear gradient to 85% over 4 minutes. The column was subsequently washed in 100% methanol for 2 minutes before being equilibrated at starting conditions. The eluate was analyzed by a Waters TQS mass spectrometer with the capillary set at 2.00 kV, desolvation temperature of 500° C, and desolvation gas flow of 900 L/hr. A multiple reaction monitoring approach was used to analyze romidepsin (541.4 > 424.2 (quantifier) and 272.4, (qualifier)) and romidepsin-d7 (548.2 > 424.4) with argon as the collision gas. Peak areas were determined in TargetLynx against a standard curve prepared in plasma. Romidepsin concentration in all nanopolymers was quantified similarly with a shorter chromatography gradient from 20% methanol to 100% methanol over 1 minute, held for 0.5 minute, before re-equilibrating for an additional 0.5 minute.

### Cell Viability Assay

Cell lines were plated at the appropriate cell densities (SUP-T1, HH, H9, FEPD, NKL and TL-1 at 100,000 cells/ml/well and SUP-M2 and FM3-29 at 50,000 cells/ml/well) in a 48-well plate. Nanoromidepsin mPEG-PDLLA PBS, Nanoromidepsin mPEG-PDLLA H_2_O, Nanoromidepsin mPEG-PLGA H_2_O, or free romidepsin were added to the cells at concentrations ranging from 0.03 nM to 30 nM with a two to three-fold serial dilution. Ghost PNP of mPEG-PDLLA and mPEG-PLGA were added to the various cell lines as a standard control at varying dilutions in order to achieve the same concentrations of the polymers as replicated in the romidepsin PNP. Cells were harvested following 60 hours (Figure 1), and 48 hours (Figure 2) of drug exposure at 37°C/5% CO2 and assayed for cell viability using CellTiter-Glo® (Promega) by following manufacturer instructions. Briefly, approximately 15 minutes after the addition of CellTiter-Glo® (at a 1:1 ratio), luminescence was read on a GloMax® Discover Microplate Reader (Promega). Luminescence was normalized to untreated control which was defined as 100% viability.

**Figure 1.**
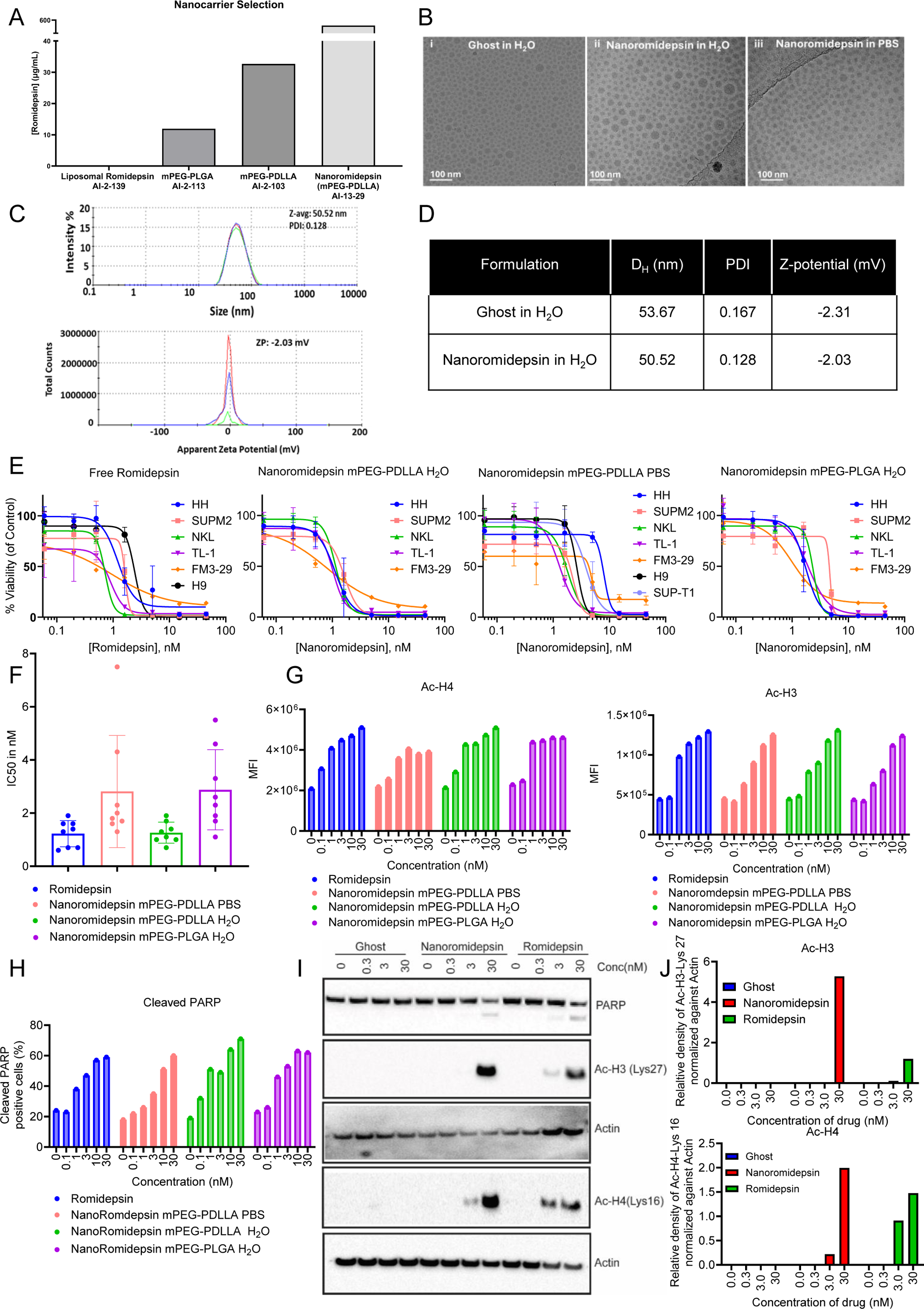
Romidepsin nanoparticle synthesis, physicochemical characterization, and drug activity analysis in PTCL cells *in vitro*. (A) Carrier selection screening. Romidepsin encapsulation quantified by LC/MS; (B) Cryo-EM of (i) Ghost in H_2_O (ii) Nanoromidepsin in H_2_O (iii) Nanoromidepsin in PBS; (C) DLS graphs (top) and Zeta-potential (bottom) spectra of Nanoromidepsin in H_2_O; (D) DLS and Zeta potential data of Nanoromidepsin ghost and Nanoromidepsin in H_2_O; (E) HH and H9 (CTCL), SUP-M2 (ALK+ ALCL) TL1 (LGL-leukemia), NKL (NK cell lymphoblastic leukemia/lymphoma), FM3-29 (melanoma) were treated with free romidepsin and different analogs of Nanoromidepsin (mPEG-PDLLA Nanoromidepsin-H_2_O, mPEG-PDLLA Nanoromidepsin-PBS and mPEG-PLGA Nanoromidepsin-H_2_O. The cytotoxicity was determined using CellTiter-Glo assay after 60 hours of treatment; (F) IC50 (nM) for romidepsin and Nanoromidepsin analogs for the 8 cell lines at 60 hours. Flow cytometry of (G) Ac-H3-lys27, and Ac-H4-lys16 (H) cleaved PARP expressing in HH cell line after 30 hours of treatment with indicated treatment of increasing concentration of free romidepsin and Nanoromidepsin. Data presented as mean ± SD; (I) Western blot analysis of Ac-H3-lys27, Ac-H4-lys16, and cleaved PARP at 24 hours after treatment with Ghost, romidepsin, and Nanoromidepsin mPEG-PDLLA H_2_O, and (J) densitometry analysis of the Western blot analysis.

**Figure 2.**
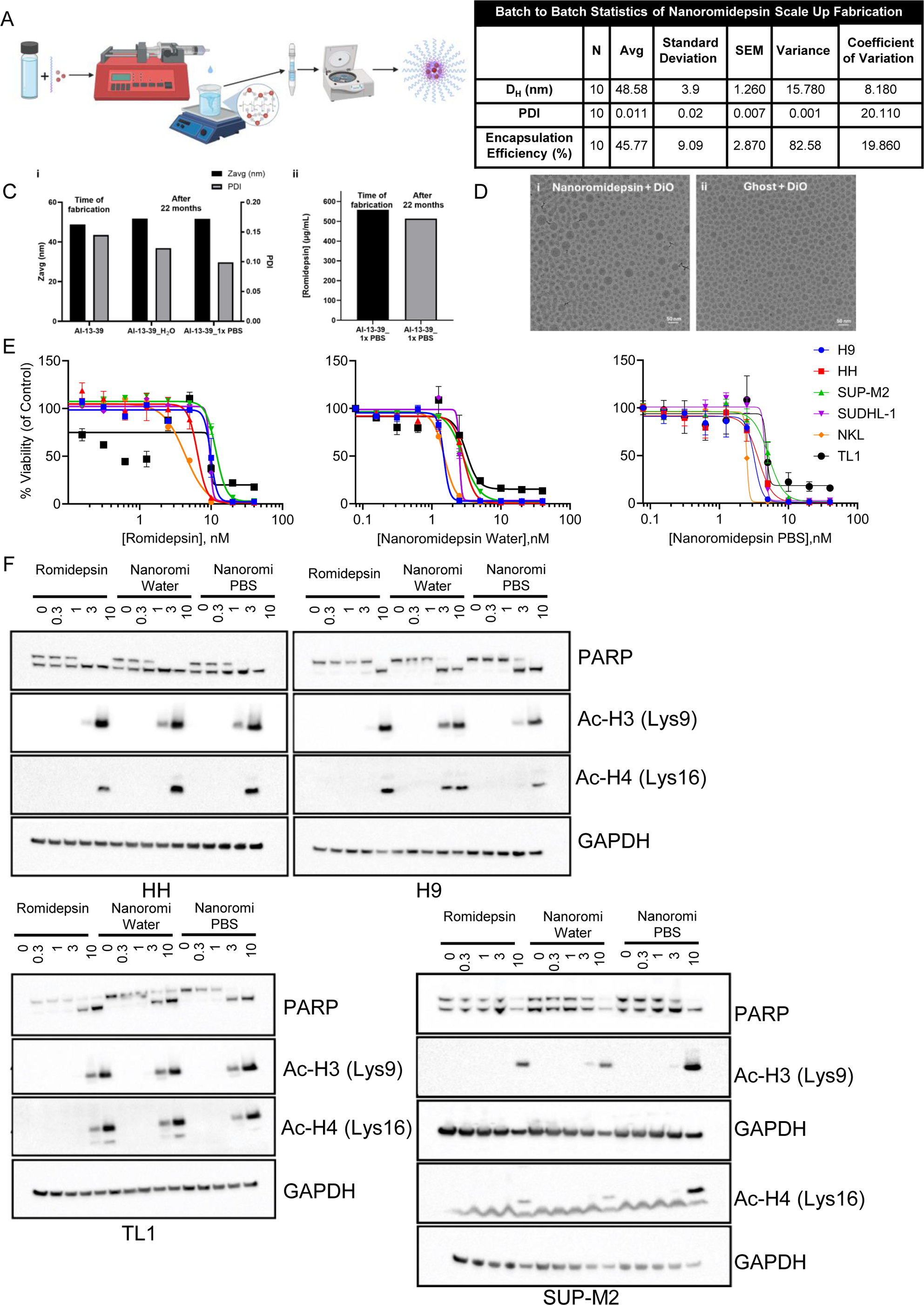
Scale-up schematic of Romidepsin polymer nanoparticle (Nanoromidepsin) synthesis, physicochemical characterization, and drug activity analysis in PTCL cells *in vitro*. (A) Schematic representation of Nanoromidepsin fabrication; (B) The statistical variability of ten, scale up (20 mL or greater) batches. Of note, these were formulated with different material stocks (polymer, Romidepsin, solvent, etc.); (C) Stability of Nanoromidepsin with respect to (i) size and PDI (ii) concentration; (D) Cryo-EM images of coloaded Nanoromidepsin with DiO and Ghost DiO; (E) HH and H9 (CTCL), SUPM2 and SUDHL-1 (ALK+ ALCL) TL1 (LGL-leukemia), NKL (NK cell lymphoblastic leukemia/lymphoma) were treated with free romidepsin and different analogs of Nanoromidepsin (mPEG-PDLLA Nanoromidepsin H_2_O and mPEG-PDLLA Nanoromidepsin PBS). The cytotoxicity was determined using CellTiter-Glo assay after 48 hours of treatment; (F) Western blot analysis of PARP, Ac-H3-K9, and Ac-H4-K16 in HH, H9, TL1 and SUP-M2 cell lines after 48 hours of treatment with indicated treatment of increasing concentration of free romidepsin and different analogs of Nanoromidepsin (NanoRomi water and NanoRomi PBS).

### Flow Cytometry

Cells were harvested after 30 hours of treatment, washed twice with PBS, fixed with 4% paraformaldehyde solution and permeabilized with 70% methanol before incubating with the relevant antibodies, including rabbit anti-HDAC1 polyclonal antibody (Proteintech; Cat no 10197-1-AP), rabbit anti-acetyl histone H3 K9/K14 polyclonal antibody (Millipore Sigma; Cat# 06-599) and rabbit anti-acetyl histone H4 polyclonal antibody (Millipore Sigma; Cat# 06-598). After 30 minutes of incubation at room temperature cells were washed twice with PBS and FITC conjugated secondary goat anti-rabbit polyclonal antibodies (Thermofisher; Cat # 31635) were added. Cells were incubated on ice for 30 minutes and then washed in PBS. After the second wash, cells were resuspended in PBS with 4% paraformaldehyde prior to signal acquisition on a flow cytometer (Attune, Thermo Fisher Scientific). Apoptosis in cell lines was assessed by the presence of cleaved poly (ADP-ribose) polymerase-1 (PARP) using a PE conjugated mouse anti-cleaved PARP (Asp214) monoclonal antibody (BD Biosciences: Cat# 552933, clone F21-852). After staining, the cells were acquired using an Attune Flow Cytometer and the data analyzed using FloJo analysis software. The Mean Fluorescence intensity (MFI) was indicative of the level of expression of each marker.

### Western Blots

Cells were incubated with the indicated concentrations of each drug and PNPs under normal growth conditions for 24 hours. Proteins from total cell lysates were resolved on 12% to 20% SDS-PAGE and transferred to nitrocellulose membranes. Membranes were blocked in phospho-buffered saline containing 0.05% Triton X-100 containing 5% skim milk powder, which were then probed overnight with specific primary antibodies. Antibodies were detected with the corresponding horseradish peroxidase–linked secondary antibodies. Blots were developed using Bio-Rad CLARITY™ and CLARITY MAX™ ECL chemiluminescent substrate detection reagents. Signal detection and imaging was captured and analyzed using a CHEMIDOC™ System (Bio-Rad) with signal quantification. The monoclonal and polyclonal antibodies used were as follows: acetylated histone H3, acetylated histone H4, vinculin, and β-actin (all from Cell Signaling Technology).

### Generation of H9-dTomato-luc Cell Line

pUltra-Chili-Luc was purchased from Addgene (*24*) (Catalog no# 48688) and was transfected into HEK293T/17 cells with lentiviral packaging plasmids VSV-G, PLP1, and PLP2 (Invitrogen) by Lipofectamine 3000 (Invitrogen) (a gift from Dr. Su-Fern Tan, University of Virginia). Viral particles were collected after 48 and 72 h post transfection. Virus particles were added to cells every 24 hours for 2 days with 7 μg/ml polybrene. Transduced cells were sorted by flow-cytometer with strongest dTomato expression and maintained in 20% FBS.

### Single and Mutiple Dose *In Vivo* Toxicity Study

5 to 7 week old female BALB/c mice (n=5) were randomly assigned to treatment cohorts based on body weight. Mice were divided into two cohorts: (i) Nanoromidepsin or (ii) free romidepsin. Mice from both cohorts were administered a single treatment of 1, 2, 3, 5, or 8 mg/kg body weight by intraperitoneal (IP) injection, and 1, 2, 3, 5, 8 or 10 mg/kg body weight by intravenous (IV) administration. The mice were monitored for 14 days post-treatment. For repeat dose maximum tolerated dose (MTD) studies, three million H9 cells expressing a fluorescent protein, Chili (dTomato-absorption max and emission max at 554 nm and 581 nm, respectively) and the bioluminescence generating protein firefly luciferase (Luc), were injected subcutaneously into the right flank of 5 to 7 week old female NSG mice (NOD-Cg-Prkdcscid Il2rgtm1Wjl/SzJ mice; The Jackson Laboratory). After the tumor reached the bioluminescent intensity (photon/s/cm2/sr) of 10^6^ or higher, the H9-dTomato-luc xenograft-containing NSG mice were treated IV as follows: (i) 2 or 3 mg/kg ghost PNP (equivalent dose), free romidepsin or Nanoromidepsin twice a week for 2 weeks; (ii) 2.5 mg/kg ghost PNP (equivalent dose), free romidepsin or Nanoromidepsin once a week for three weeks; (iii) 4 mg/kg ghost PNP (equivalent dose), free romidepsin or Nanoromidepsin once a week for three weeks, or (iv) 5 or 8mg/kg ghost (equivalent dose), free romidepsin or Nanoromidepsin once every two weeks. The mice were sacrificed the day after the last treatment. Nanoromidepsin was diluted under sterile conditions in phosphate buffered saline (PBS). Romidepsin was prepared in DMSO and then diluted with PBS under sterile condition. For single dose and repeat dose toxicity studies, weight loss and clinical score were quantitated as a function of time. Clinical signs were scored by observing activity, appearance (hair coat and eyes/nose), posture, and body condition with a maximum of 3 points ascribed to each criterion (0, normal; 1, slight deviation from normal; 2, moderate deviation from normal, 3, severe deviation from normal). Criteria leading to euthanasia included weight loss of >20% or a clinical score > 6.

### Pharmacokinetic Study

5 to 7 week old female BALB/c mice were divided into two treatment cohorts including Nanoromidepsin or free romidepsin. Each treatment cohort was further divided into two sub-cohorts depending upon the route of administration (IP or IV). Each sub-cohort (n=21) received a single treatment of one half MTD as defined from our single dose toxicity study (2.5 mg/kg body weight) of Nanoromidepsin or free romidepsin. Mice were sacrificed (n=3 per time point) at 1, 3, 6, 18, 24, 48, and 72 hours after the treatment. Blood (∼1 mL) was collected via terminal cardiac puncture under isoflurane anesthesia and collected in EDTA-coated K3EDTA tubes followed by centrifugation (2,000×g for 15 min) to isolate plasma. Plasma was placed in cryopreservation vials and preserved by snap freezing using liquid nitrogen. Blood from three untreated mice was collected at the beginning of the treatment (T=0) and used as a control. The level of romidepsin was quantified by a validated method based on reversed-phase liquid chromatography coupled to tandem mass-spectrometric detection with a standard curve derived with stock romidepsin, as described above.

Blood was collected by sub-mandibular bleeding after 1 and 24 hours following the last treatment with 4 mg/kg free romidepsin and Nanoromidepsin as described above in the repeat dose study. Plasma was collected and the romidepsin was quantified as described above. Liver and tumor were harvested, fixed in formalin for pathological analysis following H&E staining and processed for LC-MS based quantification of romidepsin.

### Biodistribution Study

To measure the biodistribution of Nanoromidepsin, NSG mice were injected with H9-dTomato-luc cells subcutaneously. The tumor growth was monitored by *in vivo* bioluminescence intensity up to twice weekly. Prior to imaging, each mouse was injected by the IP route with the bioluminescence substrate (Nanolight luciferin, 135 mg/kg body weight was given by IP with an injection volume of approximately ∼200ul). After the tumor reached a bioluminescence intensity (photon/s/cm^2^/sr) of 10^6^ or higher, tumor-bearing NSG mice were randomly assigned into two groups (n = 3) and injected intravenously with Nanoromidepsin co-loaded with DiO (fluorescent dye) or free DiO at an equivalent dose (3.7 mg/kg). Whole-body fluorescence imaging was performed at predetermined times on a cryogenically cooled Lago X (Spectral Instruments Imaging system). Three mice from each group were sacrificed after 72 hours. Tumor and all vital organs were harvested. An *ex vivo* imaging was performed of tumor and other organs.

### Survival and Efficacy Study

H9-dTomato-luc cells were injected subcutaneously into the right flank of 5 to 7-week-old female NOD Cg-Prkdcscid Il2rgtm1Wjl/SzJ mice (The Jackson Laboratory). Mice were imaged twice a week starting six days after inoculation of cells. Once tumors reached the pre-specified bioluminescence intensity (>10^6^ photon/s/cm2/sr), mice were randomized to four treatment groups of 9 mice each: (i) control group treated with normal PBS; (ii) ghost PNP; (iii) romidepsin (3.5 mg/kg), or (iv) Nanoromidepsin (3.5 mg/kg). All drugs were administered by tail vein once a week. Baseline imaging data were recorded for all mice the day before the first dose of the drug. The number of animals, study design, and treatment of animals were reviewed and approved by the Institutional Animal Care and Use Committee (IACUC) of the University of Virginia, Charlottesville. *In vivo* BLI analysis was conducted on a cryogenically cooled Lago X (Spectral Instruments Imaging system). A second efficacy/survival study was performed using similar methods with 4 different treatment groups of 9 animals each, including: (i) control group treated with normal PBS; (ii) ghost PNP; (iii) romidepsin (4 mg/kg) and (iv) Nanoromidepsin (4 mg/kg).

### *Ex vivo* Cytotoxicity and Flow Cytometry Study

Primary LGL leukemia cells were collected from patients (LGL Leukemia Registry at University of Virginia) who met the criteria of T-LGL (CD3+/CD8+) leukemia with increased numbers of LGL cells in the peripheral blood. Peripheral blood samples were obtained and informed consent signed for all LGL leukemia patients according to protocols approved by the Institutional Review Board of University of Virginia School of Medicine. PBMCs were isolated by Ficoll-Paque gradient separation and viably cryopreserved (*25*). Samples with viability > 90% were then used for further downstream analysis. PBMC were seeded with 100,000 cells/well in a 96 well plate and treated with indicated doses of ghost PNP, romidepsin and NanoRomidepsin. Cytotoxicity assay was performed using Cell Titer Glo as described above.

Flow cytometry analysis was performed as described earlier (*26, 27*). Briefly, PBMCs from the same patients used in the cytotoxicity assay were treated with the indicated doses of ghost PNP, romidepsin and Nanoromidepsin for 48 hours. The PBMCs were stained with Live/Dead near-infrared viability dye. Cells were then fixed in paraformaldehyde (1.6%) and stained for surface markers using anti– CD3BV605(clone OKT3), anti-CD8 BV421(clone SK1), and anti– CD57 BB515 (clone NK-1), for overnight at 4°C. PBMCs were washed and permeabilized using saponin and stained with anti-cleaved PARP-PE (clone F21-852). Flow cytometry data analysis was performed using FlowJo software.

### Statistical Analysis

Results are presented as the mean ± SD, unless indicated otherwise. Statistical significance was determined by 1-way ANOVA or 2-tailed Student’s t test or log rank test, unless specified otherwise, using GraphPad Prism software, and a p-value of less than 0.05 was considered statistically significant.

## Results

### Engineering of Nanoromidepsin and DiO Loaded Polymer Nanoparticles (PNP)

Several different NPs of romidepsin were synthesized using generally regarded as safe (GRAS) amphiphilic di-block copolymers and FDA approved lipids for liposomes. Liposomes did not achieve romidepsin encapsulation and were not pursued further. PNPs were synthesized using mPEG-PDLLA and mPEG-PLGA, and the surfactant poloxamer-188 using a solvent displacement or nanoprecipitation technique as described above. LC/MS confirmed an average romidepsin concentration in liposomes and polymer nanoparticles of <0.1ug/mL and >500ug/mL respectively. (Figure 1A). mPEG-PDLLA nanoparticles exhibited higher drug concentrations of approximately 540 µg/mL with an average EE of 48%. Cryo-EM (Figure 1B) revealed that both ghost and romidepsin loaded PNPs exhibited uniform spherical morphology, homogeneous size of 35 nm with no agglomeration or agglomeration. DLS (Figure 1C-D) revealed a unimodal distribution of the particles with an average size of 46.25 nm and a PDI of 0.145.

The concentration-response relationship for each of the PNP was determined and compared to free romidepsin across a panel of T-cell lymphoma and LGL leukemia cell lines, as well as a melanoma cell line to understand activity in a solid tumor malignancy (Figures 1E). The IC_50_ for each PNP is shown in Figure 1F. Ghost PNPs were diluted to match the volume of the polymer shell in the Nanoromidepsin derivatives (Figure S1), and consistently revealed minimal to no toxicity in the *in vitro* assays. All three PNPs of romidepsin inhibited cell growth of all cancer cell lines in a concentration dependent manner (Figure 1E), though the IC_50_ values for the three different PNPs varied across the cell lines tested. At 60 hours, most cell lines were consistently sensitive to Nanoromidepsin mPEG-PDLLA H_2_O (IC_50_= 0.7-1.9 nM) which was very similar to romidepsin (IC_50_ =0.6-1.9 nM) (Figure 1F). Both Nanoromidepsin mPEG-PDLLA PBS (IC_50_= 1.3-7.5 nM) and Nanoromidepsin mPEG-PLGA H_2_O (IC_50_=1.1-5.5) were slightly less potent compared to romidepsin and Nanoromidepsin mPEG-PDLLA H_2_O. There was no inhibition of growth in any cell line with the corresponding ghost PNP. (Figure S1). Flow cytometry and western blotting demonstrated that treatment with all three romidepsin PNPs induced apoptosis similar to romidepsin as shown by an increase in the expression of cleaved PARP (Figure 1H and 1I). However, Nanoromidepsin mPEG-PDLLA PBS and Nanoromidepsin mPEG-PDLLA H_2_O demonstrated higher PARP cleavage compared to romidepsin and Nanoromidepsin mPEG-PLGA H_2_O.

Levels of H3 and H4 acetylation were measured following treatment with increasing concentrations of romidepsin and the three PNPs of romidepsin. A concentration dependent increase in acetylation of both H3 and H4 was observed by flow cytometry when cancer cells were treated with either romidepsin or one of the three romidepsin PNPs (Figure 1G). Although a similar degree of increased histone acetylation was observed after treatment with all three PNPs, Nanoromidepsin mPEG-PDLLA H_2_O was comparable to free romidepsin in its pattern of histone acetylation. Given that among the three PNPs, Nanoromidepsin mPEG-PDLLA H_2_O exhibited the lowest IC_50_ and comparable histone acetylation and PARP cleavage compared to free romidepsin, we further validated the flow cytometry data with western blotting analysis with Nanoromidepsin mPEG-PDLLA H_2_O. The levels of acetylation of H3 lysine 27 and H4 lysine 16 were measured following treatment with increasing concentrations of romidepsin and Nanoromidepsin mPEG-PDLLA H_2_O. Western blot analysis demonstrated an increase in H3/H4 acetylation following treatment with 0.3 to 30 nM of romidepsin or Nanoromidepsin mPEG-PDLLA H_2_O at 24 hours (Figure 1I and IJ). The increased acetylation of both H3 and H4 proteins was higher in cells treated with Nanoromidepsin compared to romidepsin.

Between the mPEG-PLGA and mPEG-PDLLA Nanoromidepsin, the PDLLA Nanoromidepsin showed the best physicochemical properties (size, PDI, and encapsulation efficiency) and biological activity, prompting further optimization, scale up, and characterization (data not shown).

### Scale-up of Romidepsin PNP Synthesis, Physicochemical Characterization, and *in vitro* Activity

The scale up workflow and resulting PNP physicochemical properties are shown in Fig 2A-B. Several key factors influencing the batch production of romidepsin PNP were identified and optimized for bulk synthesis. These factors include: the polymer type, drug-to-polymer ratio, drug-to-surfactant ratios, diffusion coefficients of solvents, solvent-to-anti-solvent ratio, rate of addition of reagents and processing (evaporation, centrifugation, and reconstitution) of nanoparticles (Figure 2A-2B). These factors play a critical role in the efficiency of drug encapsulation, scalability, nanoparticle characteristics such as size, PDI, drug release, and the nature of discrete biological interactions. The Nanoromidepsin PDLLA H_2_O revealed particle stability for 22 months at 4°C (Figure 2C). Co-loading of romidepsin with DiO produced particles of similar size, indicating this method can potentially accommodate efficient co-packaging of multiple drugs. (Figure 2D).

To verify that the scaled-up version of Nanoromidepsin PDLLA H_2_O and Nanoromidepsin PDLLA PBS retained similar cell-killing properties without compromising the mechanism of action, we repeated cytotoxicity studies to assess the cytotoxicity of the PNPs across the panel of PTCL and LGL leukemia cell lines (HH, H9, SUP-M2, SUDHL1, TL1, and NKL). All six TCL cell lines showed varying degrees of sensitivity to romidepsin and Nanoromidepsin PNPs after 48 hours of treatment (Figure 2E). The IC_50_ of Nanoromidepsin PNPs were ∼2-fold lower (IC_50_ between 1.5-5 nM) compared to free romidepsin (IC_50_ value 4-10 nM) (Figure S2). Next, we performed western blot analysis on four different TCL cell lines to compare the effect of the scaled up Nanoromidepsin PDLLA H_2_O and Nanoromidepsin PDLLA PBS PNPs on histone acetylation (Figure 2F). After 48 hours of treatment, Nanoromidepsin PNPs demonstrated superior or comparable H3/H4 acetylation compared to free romidepsin across all cell lines studied. HH, H9, and SUPM2 cell lines demonstrated a substantially greater increase in histone acetylation after treatment with Nanoromidepsin PNPs compared to free romidepsin consistent with the data in Fig. 2E, affirming a relatively lower IC_50_ value for Nanoromidepsin PNPs (Figure 2F).

### Nanoromidepsin Exhibited Superior Cytotoxicity Against Primary LGL Leukemia Samples

The anti-proliferative effect of free romidepsin and Nanoromidepsin were compared on PBMC isolated from LGL leukemia patients. LGL-leukemia is a chronic mature lymphoproliferative disorder that is distributed into two subtypes: T-LGL leukemia and NK-LGL leukemia (*28*). T-LGL leukemia typically exhibits a CD3+, TCR αβ+, CD4-, CD5dim, CD8+, CD16+, CD27-, CD28+, CD45RO-, CD45RA+, and CD57+ phenotype, which represents a constitutively activated T-cell phenotype (*29–31*). Nanoromidepsin demonstrated superior cytotoxicity in TL1 (a T-cell LGL) and NKL (a NK cell LGL) cell lines, exhibiting a lower IC_50_ compared to free romidepsin (Figures 2A and B). An *ex vivo* cytotoxicity assay performed on PBMC from the LGL-leukemia patients demonstrated that Nanoromidepsin produced a statistically significant improvement in potency, with an IC_50_ of 3.1 ± 1.7 nM, compared to free romidepsin, which exhibited an IC_50_ of 9.06 ±5.7 nM. (p=0.0057) (Figure 3A and 3B.). As whole PBMC samples also contain a small proportion of non-leukemic cells, we designed a multi-color flow cytometry-based functional assay (*27*) to quantify apoptosis in CD3+CD8+CD57+ or CD3+CD8+CD57-cell populations of LGL-leukemia patients (Figure 3C). PBMCs from healthy individuals were used as control. These data revealed that the percentage of CD3+CD8+CD57- and CD3+CD8+CD57+ cells positive for the cleaved PARP apoptosis marker was similar for Nanoromidepsin and free romidepsin treated PBMC samples, though the percentage of dead cells (viability dye+) in CD3+CD8+CD57+ and CD3+CD8+CD57-cells was comparatively higher in the Nanoromidepsin treated PBMC samples although the results were not statistically significant (p< 0.59 and 0.46 respectively) (Figure 3C and 3D).

**Figure 3.**
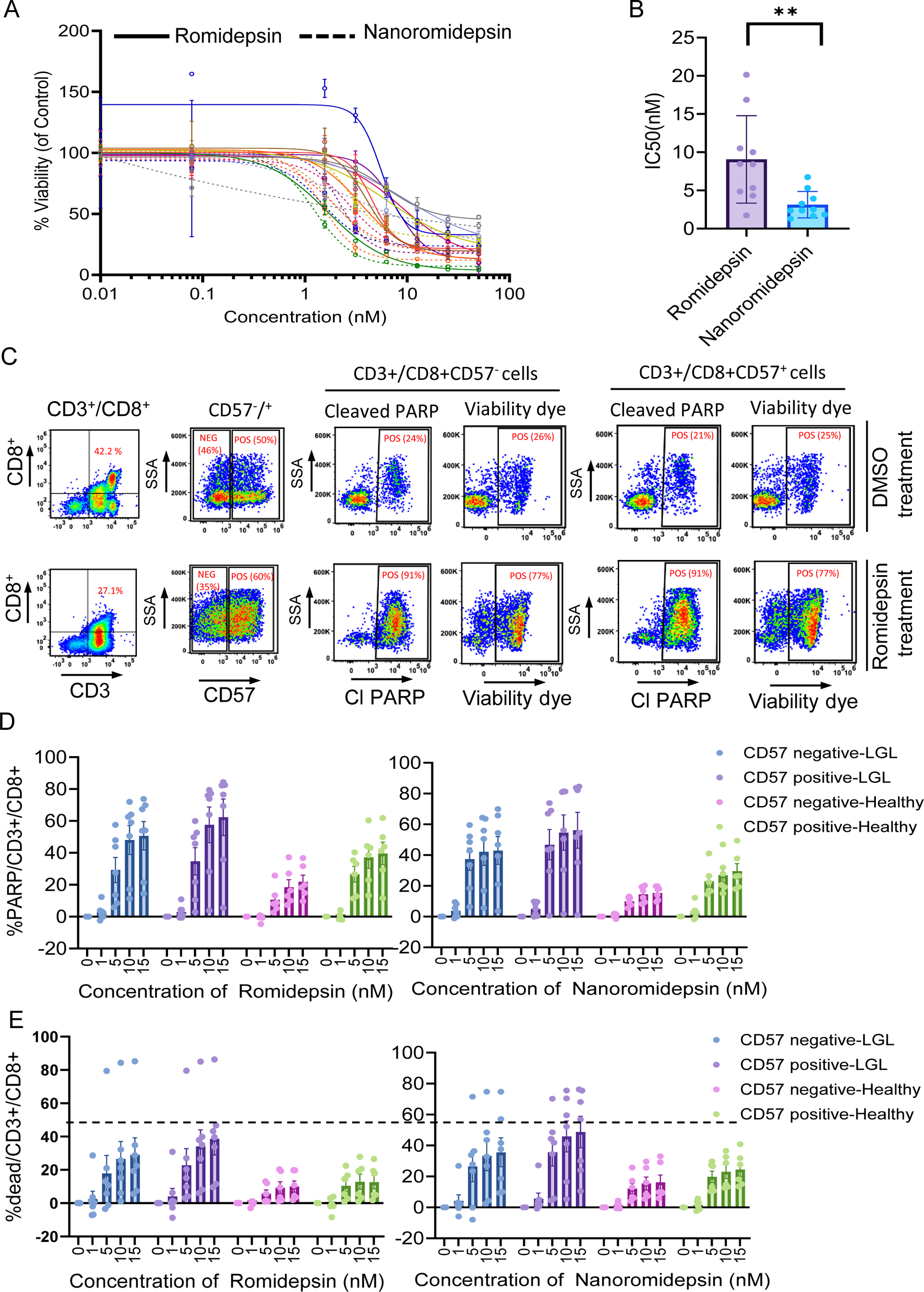
Effect of Nanoromidepsin on Primary LGL Leukemia patient PBMC samples. (A) Freshly frozen PBMCs from LGL leukemia patients were treated with indicated doses of romidepsin (solid line) or Nanoromidepsin (dotted line) for 48 hours. The cytotoxicity was determined using CellTiter-Glo assay after 48 hours of treatment; (B) IC50 (nM) for romidepsin and Nanoromidepsin for 10 LGL leukemia patients at 48 hours; (C) PBMCs from patients with LGL leukemia and healthy donor control were screened by flow cytometry. The lymphocyte and singlet cell gating were performed as described earlier. The CD3+/CD8+/CD57+/− cells were gated from singlet lymphocyte population as indicated. The cleaved PARP or viability dye staining was analyzed in CD3+/CD8+/CD57+ or CD3+/CD8+/CD57^−^ cells as indicated. The flow images were generated from a representative LGL patient (PT #03) PBMC sample treated with DMSO or romidepsin (10 nM). Ghost or Nanoromidepsin treated samples were similarly analyzed; (D) Cleaved PARP (apoptosis) and (E) live-dead dye staining (cell viability) after the incubation with free romidepsin and Nanoromidepsin for 48 hours. Data presented as percentage CD3^+^/CD8^+^/ CD57^+^ (more differentiated LGL) or CD3^+^/CD8^+^/ CD57^−^ (less differentiated LGL) cells positive for cleaved PARP or live-dead dye staining. The data presented after subtracting spontaneous apoptosis or cell viability values from the DMSO-treated controls.

### Nanoromidepsin Demonstrates Superior Pharmacokinetic Parameters and Biodistribution Compared to Free Romidepsin

To evaluate the pharmacokinetic profile of the Nanoromidepsin PDLLA PBS (hereafter referred to as Nanoromidepsin), BALB/c mice were injected with the indicated drugs by IV or IP administration. The plasma concentration of romidepsin was quantified using LC-MS/MS. Irrespective of the route of administration, the plasma romidepsin concentration of free romidepsin rapidly declined after 6 hours. (Figure 4A). In contrast, Nanoromidepsin exhibited a higher area under the curve (AUC) of exposure 48 hours post-treatment, irrespective of the route of administration. After IV administration, Nanoromidepsin attained peak drug concentrations (T_max_) at 6 hours, while free romidepsin attained a T_max_ of 3 hours. The peak concentration (C_max_) and AUC for Nanoromidepsin by the IV route were 10-fold and 25-fold higher compared to free romidepsin respectively (Table 1). The pharmacokinetic (PK) analyses also indicated that the clearance of romidepsin from the plasma by the IV route of administration was faster compared to the IP route of administration. The peak concentration of romidepsin achieved after IP administration of Nanoromidepsin and romidepsin were 804 nM and 218 nM, respectively. After IV administration, the peak concentration of Nanoromidepsin and free romidepsin were 425 nM and 38 nM, respectively. Based on the *in vitro* data across the TCL cell lines studied, the IC_50_ of Nanoromidepsin PDLLA was around 2 to 8 nM. These data suggest that Nanoromidepsin can achieve a concentration 80-400-fold greater than the IC_50_ of romidepsin with a dose that was only one-half of the MTD dose of Nanoromidepsin (described below in Figure 5A).

**Figure 4.**
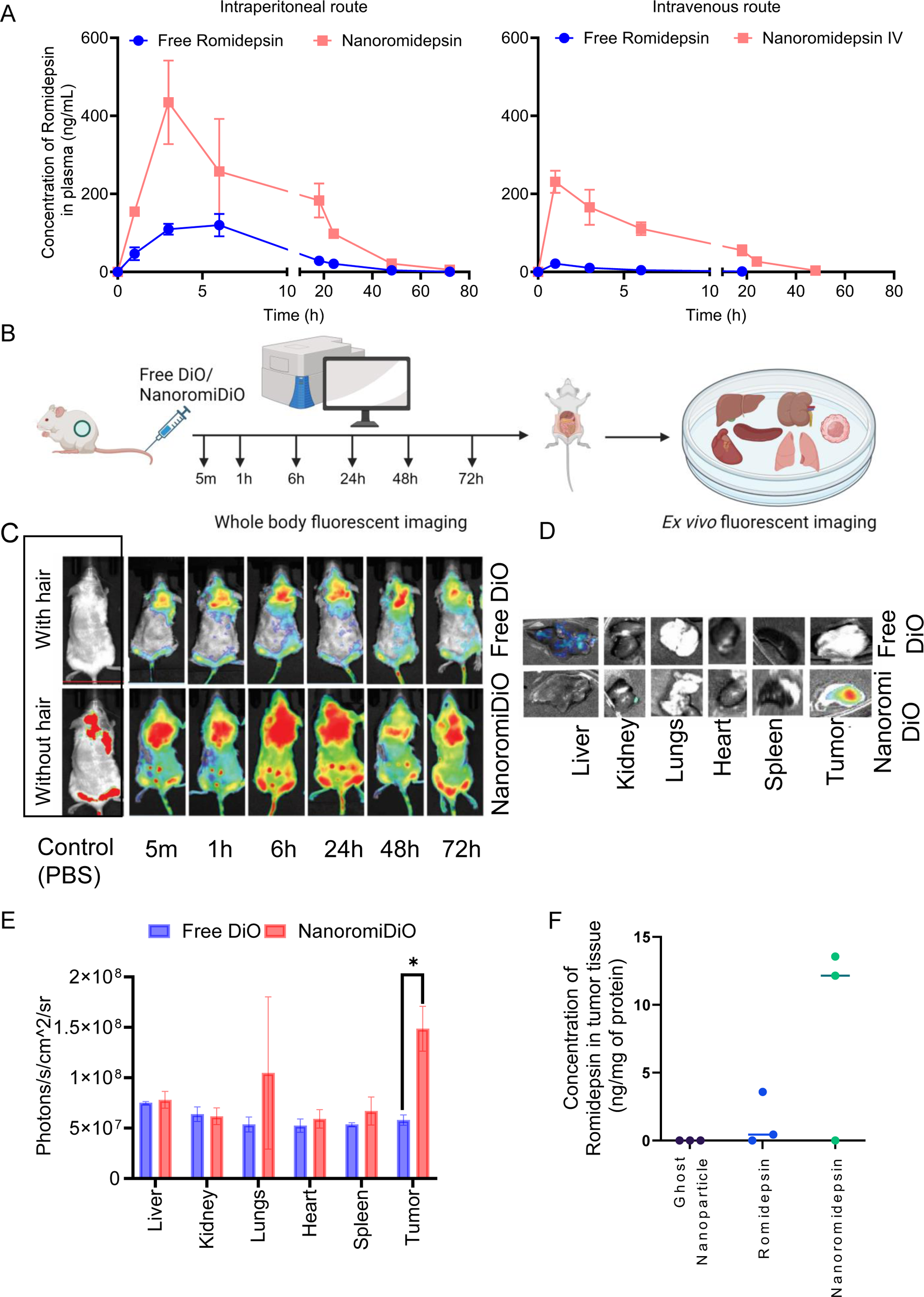
Pharmacokinetics and tissue distribution of Nanoromidepsin *in vivo*. (A) Plasma concentration-time dependence plot of romidepsin concentration in plasma after intraperitoneal or intravenous administration of a single treatment with free romidepsin or Nanoromidepsin; (B) Diagram representing experimental time-points associated with Nanoromidepsin co-loaded with a fluorescent dye DiO or free DiO administration and fluorescent images evaluation, as well as organs collection; (C) Fluorescence images of H9-dTomato-luc tumor-bearing mice taken at different time points after intravenous injection of free DiO or DiO and romidepsin encapsulated nanoparticle (NanoromiDiO); (D) *Ex vivo* fluorescence images and (E) corresponding optical intensity of tumor and major organs (tumor, liver, spleen, kidney, heart, and lung, respectively) dissected at 72 h post-injection. Statistical significance was determined by using student t test (Mann-Whitney) where *, p<0.05, **< p<0.01, ***, p<0.001; (F) Mice bearing H9-dtomato-luc xenograft were treated with 4 mg/kg romidepsin and Nanoromidepsin. After 24h, tumors (n=3) were collected for LC-MS based quantification of romidepsin in tumor tissue.

**Figure 5.**
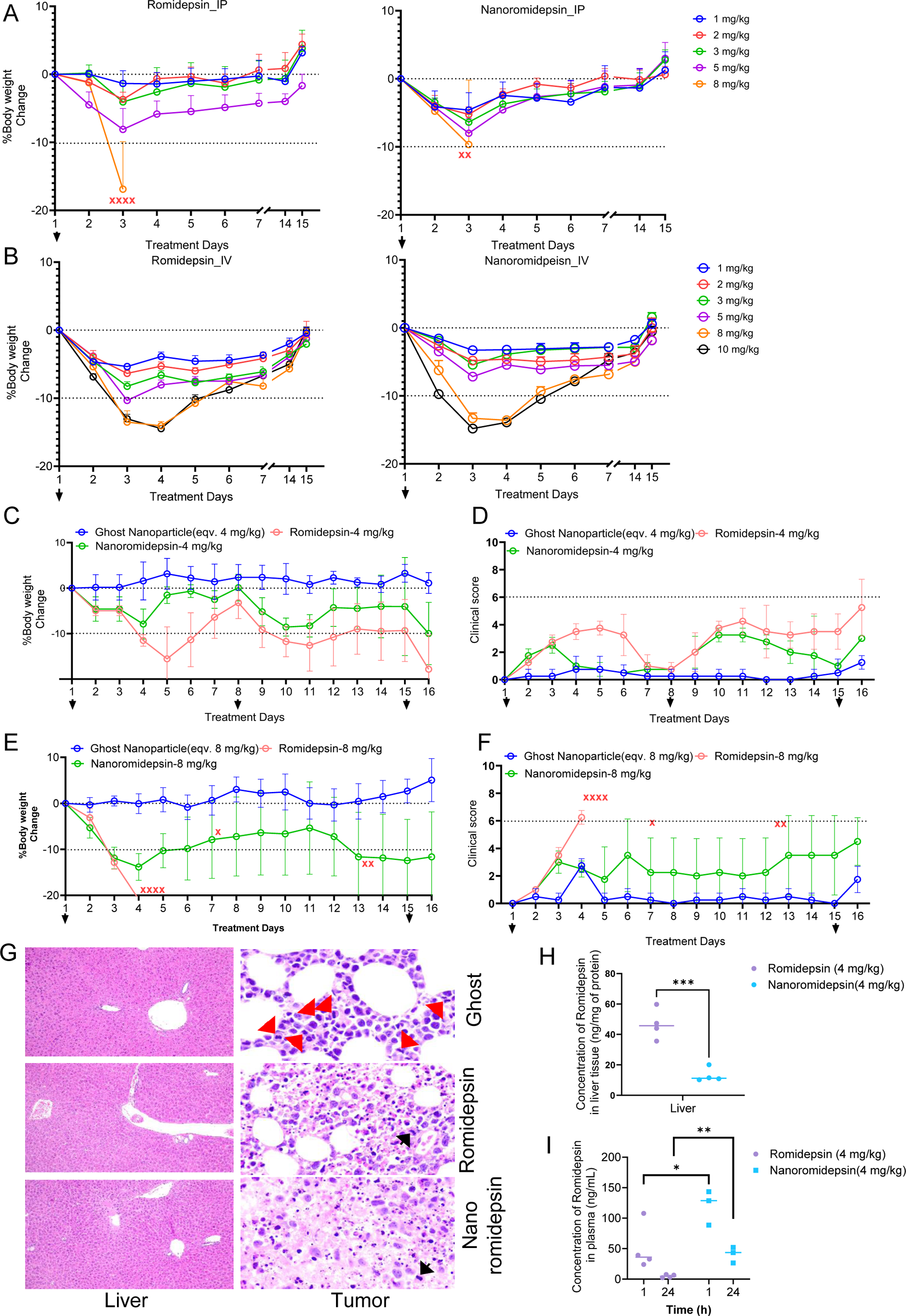
Tolerability of free romidepsin and Nanoromidepsin *in vivo*. BALB/c mice were administered a single treatment of indicated doses of romidepsin and Nanoromidepsin by (A) intraperitoneal (B) intravenous route of administration. Tolerability of various doses of free romidepsin and Nanoromidepsin in BALB/c mice was assessed by monitoring body weight and overall health conditions after a single treatment. X represents dead mice; (C) Mice were administered a 4 mg/kg treatment of romidepsin or Nanoromidepsin by intravenous route of administration for arrow indicated days (1, 8 and 15 days) in H9-dTmato-luc xenograft bearing NSG mice. Depicted are the percentage of body weight changes as a percentage of starting weights with SEM (D) clinical score. Mice were administered an 8 mg/kg treatment of romidepsin or Nanoromidepsin by intravenous route of administration for arrow indicated days (1 and 15 days) in H9-dTmato-luc xenograft bearing NSG mice. Depicted are the (E) percentage of body weight changes as a percentage of starting weights with SEM and (F) clinical score as defined in Materials and Methods; (G) The hepatic parenchyma from ghost, romidepsin, and Nanoromidepsin treated mice display normal microarchitecture, without evidence of drug-induced liver injury (original magnifications X200; H&E stain). Soft tissue-based tumors from ghost-treated mice contain sheet-like infiltrates of large atypical lymphocytes with pleomorphic nuclei, distinct nucleoli and amphophilic cytoplasm; mitotic activity is brisk (original magnification X1000; H&E stain). Romidepsin and Nanoromidepsin-treated tumors are associated with varying degrees of treatment-related necrosis (original magnifications X1000; H&E stain). Red arrows mark mitotic figures and black arrow indicates necrosis/apoptosis; (H) LC-MS based quantification of liver (I) LC-MS based quantification of plasma collected from 4C experiment after 1 and 24 hours. Statistical significance was determined by using student t test (Mann-Whitney) where *, p<0.05, **< p<0.01, ***, p<0.001.

**Table 1.** Pharmacokinetic parameters of romidepsin and Nanoromidepsin after IP and IV route of administration.

| Route of administration |  | Intraperitoneal (IP) |  |  | Intravenous (IV) |  |  |
| --- | --- | --- | --- | --- | --- | --- | --- |
| Parameter | Unit | Free Romidepsin | Nano romidepsin | Fold change (Nano/Free) | Free Romidepsin | Nano romidepsin | Fold change (Nano/Free) |
| T1/2 | h | 9.8 | 11.6 | 1.2 | 5.2 | 7.6 | 1.5 |
| Tmax | h | 6.0 | 3.00 | 2 | 1.00 | 1.00 |  |
| Cmax | ng/ml | 119.9 | 434.7 | 3.6 | 21.3 | 231.0 | 10.8 |
| AUC 0-t | ng/ml*h | 1918.9 | 6939.9 | 3.6 | 99.2 | 2532.1 | 25.5 |

T1/2- Half life; Tmax-time to maximum plasma concentration; Cmax- maximum plasma concentration; AUC 0-t- the area under the curve up to the last quantifiable time-point

To characterize the biodistribution of Nanoromidepsin, time-dependent tissue and tumor uptake studies were performed. Mice engrafted with the H9-dTomato-luc xenograft were administered Nanoromidepsin co-encapsulated with DiO fluorescent dye by the IV route of administration, in parallel with a control group administered free DiO. Whole body fluorescent images were taken at different time points as illustrated in (Figure 4B). The organs were harvested from the mice after 72 hours and *ex vivo* fluorescent images were taken. The whole-body imaging of the mice showed that the fluorescence signal of Nanoromidepsin-DiO treated mice was greater compared to the free DiO treated mice even 5 minutes post-administration and throughout the time course (Figure 4C). *Ex vivo* analysis of the organs showed that Nanoromidepsin selectively accumulated in the tumor at 72 hours post-administration, with modest uptake in the liver, which was observed only in free DiO treated mice. (Figure 4D). The quantification of fluorescent signal in all harvested organs showed a significant (p<0.05) accumulation of Nanoromidepsin in the tumor compared to the free DiO (Figure 4E). In a similar approach, H9-dTomato-luc engrafted mice were injected with 4 mg/kg romidepsin and Nanoromidepsin. Quantitation of romidepsin by LC-MS/ MS in the tumor 24 hours post-administration of Nanoromidepsin and free romidepsin (IV) revealed an intertumoral concentration of romidepsin in the free romidepsin and Nanoromidepsin treated groups of 1.34 and 8.57 ng/mg of protein respectively, suggesting a substantially greater accumulation of drug in animal treated with the Nanoromidepsin. (Figure 4F).

### Nanoromidepsin Exhibited Superior Tolerability Compared to Free Romidepsin *In Vivo*

The safety and tolerability of Nanoromidepsin was determined in a single dose toxicity study in female BALB/c mice. Mice were treated with incremental doses of Nanoromidepsin or free romidepsin (IP and IV route) to identify the MTD after single treatment. Changes in body weight and clinical score were assessed as a function of time and dose. While mice in both treatment cohorts experienced weight loss post treatment, the weight returned to pre-treatment levels in most animals after 15 days (Figures 5A-B and Figure S3). Mice treated with 8 mg/kg of either free romidepsin or Nanoromidepsin by the IP route met criteria for euthanasia three days post-treatment. At this dose level, about 80% of the mice (4 of 5 mice) treated with romidepsin were found dead three days post-treatment, while only about 40% Nanoromidepsin treated mice (2 of 5 mice) were found dead on the same day. The MTD for both drugs when administered by the IP route was 5 mg/kg. By the IV route of administration, the highest dose studied with each drug was 10 mg/kg. Mice lost approximately 15% body weight within three days after treatment with both 10 mg/kg free romidepsin and Nanoromidepsin, although all mice in both treatment groups recovered after 15 days. Escalation beyond 10 mg/kg was technically not feasible given the volume of the intravenous dose required. Thus, the MTD for both drugs when administered by IV was determined to be 10 mg/kg.

Although the AUC and C_max_ of Nanoromidepsin were considerably higher when drug was administered intraperitoneally compared to the intravenous route of administration, a study in H9 xenograft engrafted NSG mice confirmed that the IP route for Nanoromidepsin exhibited unacceptably high toxicity (Figure S4). These findings were consistent with the literature suggesting that many nanoparticles cannot be administered safely by IP owing to the association with peritonitis likely due to the physical features of the particle (*32*). For these reasons, all *in vivo* studies hereafter exclusively utilized the intravenous route.

Multiple dose and schedule studies were conducted to assess repeat dosing toxicity in H9-dTomato-luc xenograft-containing mice (Table 2 and Figure S5 and S6). The optimum dose and schedule for Nanoromidepsin was identified to be 4 mg/kg weekly for three weeks followed by a one-week rest (Figure 5C-D). Repeat dosing studies revealed that free romidepsin produced a higher degree of weight loss (>10%) and clinical score (>3) compared to Nanoromidepsin, demonstrating the superior tolerability of the PNP. Free romidepsin at a dose of 8 mg/kg in H9-dTomato-luc xenograft-containing NSG mice demonstrated acute toxicity leading to death of all mice (thus LD50 is significantly less than 8 mg/kg) within four days, while 8 mg/kg Nanoromidepsin was lethal in only 50% of mice, representing the LD50 of Nanoromidepsin. (Figure 5E and 5F).

**Table 2.** Description of different dosing schedules used to define MTD in repeated dose toxicity study.

| Dose | Schedule | Figure |
| --- | --- | --- |
| 2.5 mg/kg | Weekly x 3 | Figure S6 |
| 4 mg/kg | Weekly x 3 | Figure 5 |
| 2 mg/kg | Twice weekly x 2 | Figure S5 |
| 3 mg/kg | Twice weekly x 2 | Figure S5 |
| 5 mg/kg | Once in every 2 weeks | Figure S6 |
| 8 mg/kg | Once in every 2 weeks | Figure 5 |

To assess organ-specific toxicity, mice were treated with 4 mg/kg of free romidepsin, ghost PNP or Nanoromidepsin as detailed in Figure 5C and 5D (same experiment). Twenty-four hours after the last treatment, liver and tumor were harvested and assessed for pathological findings (Figure 5G). Liver sections from all mouse cohorts showed normal microarchitecture without any indication of inflammation or necrosis. Tumor sections from the mice treated with the ghost PNP revealed sheet-like infiltrates of large atypical lymphocytes with pleomorphic nuclei, distinct nucleoli and amphophilic cytoplasm, consistent with viable tumor. The romidepsin and Nanoromidepsin treated tumor sections showed varying degrees of treatment related necrosis, with no substantial difference in the histopathology between the two treatment groups. Although there were no signs of drug induced toxicity in the liver sections of either treatment cohort, the LC-MS confirmed that the concentrations of romidepsin in the liver tissue after the administration of free and Nanoromidepsin were 46.68 and 13.18 ng/mg of protein, respectively (p<0.0009) (Figure 5H). The mean plasma concentrations of romidepsin after 1 hour and 24 hours following three consecutive treatments of romidepsin (weekly doses for three weeks) were 51 and 4.9 ng/mL (Figure 5I). These data suggest a rapid decline in mean plasma concentration 24 hours after the third dose, implying a rapid clearance of free romidepsin from the blood. In contrast, the mean plasma concentrations of romidepsin in the plasma collected at 1 and 24 hours at the same dose of Nanoromidepsin were 120.3 and 40.7 ng/mL, (2.3 and 8.3-fold greater than the free drug).

### Nanoromidepsin Showed Superior Activity and a Survival Advantage in Murine Xenograft Models

To determine efficacy and differences in overall survival (OS), dTomato-luc expressing H9 cells were engrafted subcutaneously into the right flank of NSG mice. Mice were treated when the tumor luminescence reached 10^6^ bioluminescence intensity (BLI; p/s/cm2/sr), usually 6-7 days after the engraftment. Mice were treated with 3.5 mg/kg weekly for 3 weeks with free romidepsin or Nanoromidepsin (Figure 6A). After three treatments, the cohort receiving free romidepsin exhibited moderate antitumor activity with tumor growth inhibition of 54% and 57% compared to the vehicle and ghost PNP cohorts respectively (p=0.0315 vs vehicle; p=0.04 vs ghost PNP). Treatment with Nanoromidepsin showed inhibition of 90% and 91% compared to the vehicle and ghost PNP cohorts respectively (p=0.0003 vs vehicle; p=0.0019 vs ghost PNP). While there was no statistically significant difference in the growth delay observed between romidepsin and Nanoromidepsin (p=0.6665), Nanoromidepsin showed greater tumor reduction compared to free romidepsin after 3 weeks of treatment (Figure 6B). The tumor BLI signal was reduced one week after the first treatment which held constant for the next three weeks for both treatment cohorts (Figure 6B, D and E). Interestingly, the Nanoromidepsin treated cohort showed delayed tumor growth compared to romidepsin until the third week of treatment. However, mice treated with Nanoromidepsin or free romidepsin did not show any statistically significant improvement in survival benefit at this dose or schedule due to cytokinetic failures, which was attributed in part to the fact mice had to receive a lower dose of drug and only one cycle of therapy (Figure 6C).

**Figure 6.**
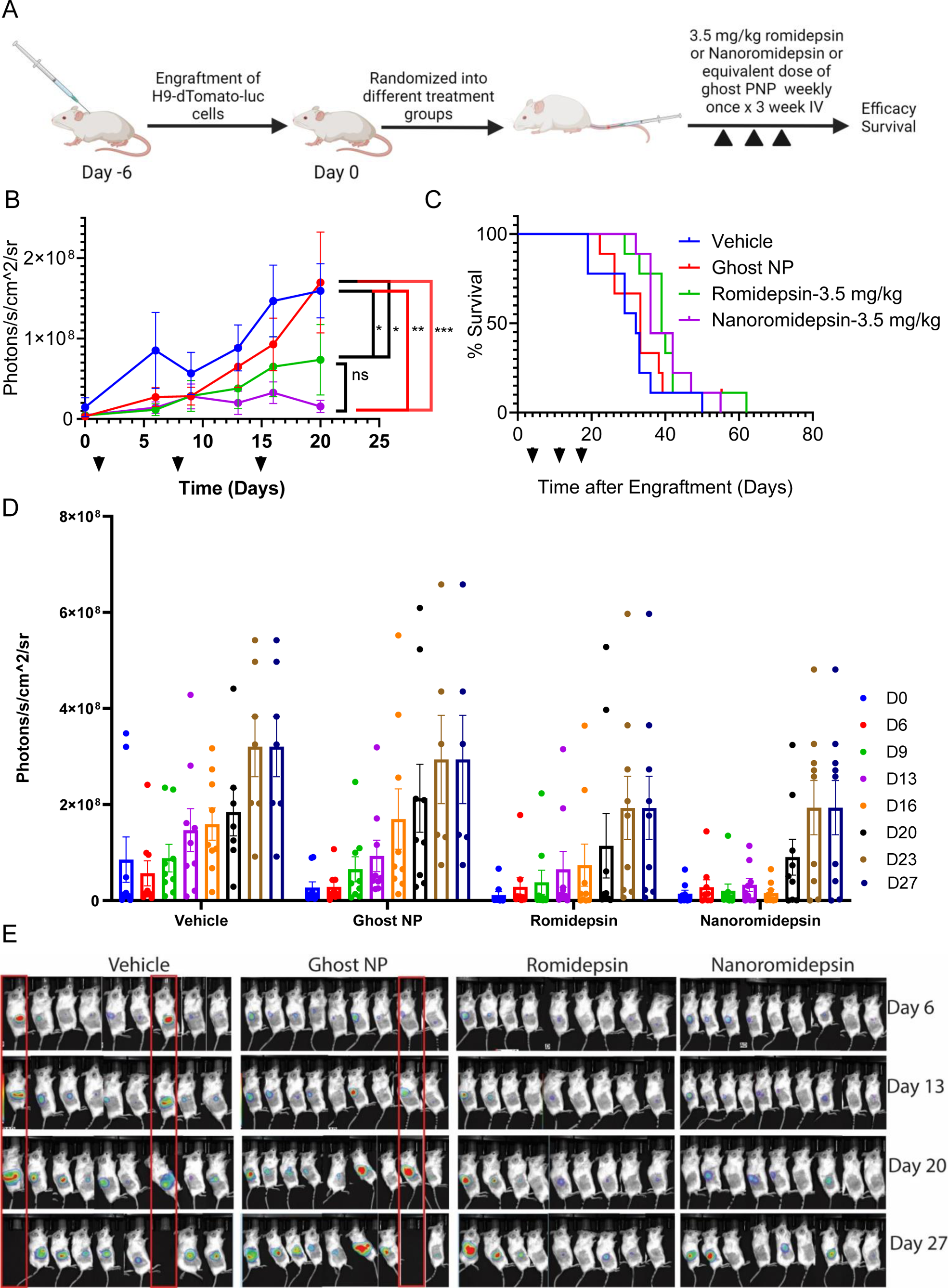
Nanoromidepsin showed superior activity but similar survival rate compared to free romidepsin in TCL xenograft bearing NSG mice. (A) Diagram representing the inoculation and dosing schedule of free romidepsin and Nanoromidepsin in H9-dtomato-luc xenograft bearing mice; (B) and (D) Region-of-interest analysis of BLIs (readout for tumor growth) from different treatment groups were recorded at various time points over the course of 8 weeks. Statistical significance was determined by using student t test (Mann-Whitney) where *, p<0.05, **< p<0.01, ***, p<0.0001; (C) Survival curves for romidepsin–treated, Nanoromidepsin-treated, ghost nanoparticle and control mice (*n* = 9 per group). The arrows indicate treatment days. Statistical significance was determined by using log rank test where *, p<0.05, **< p<0.01, ***, p<0.0001; (E) Whole-body bioluminescence images of H9-dTomato-luc xenograft–bearing mice taken at the indicated day. Red box indicates dead mouse.

Given the issue of cytokinetic failure, we explored further modifying the dose and schedule, administering both drugs at a higher dose of 4 mg/kg weekly for four consecutive weeks, followed by a two-week break (Figure 7A). Significant toxicity was noted after one treatment with free romidepsin. As shown in Figures 7B and C, a consistent increase in the BL1 intensity which is proportional to tumor growth was observed in the PBS, ghost PNP and free romidepsin treated mice cohort until day 24. However, a growth delay was observed in Nanoromidepsin treated mice in the same time frame. Moreover, as shown Figure 7C, 33% of mice died after three weeks of treatment with free romidepsin, while treatment with Nanoromidepsin resulted no deaths. Consistent with the bioluminescence imaging data, Nanoromidepsin treatment resulted in a highly statistically significant prolongation in OS compared to the free romidepsin treated mice. The overall survival in the control, ghost PNP and romidepsin treated mice was 38 days (for all three groups). In contrast, the median OS of Nanoromidepsin administered mice was 53 days (p<0.001), which was highly statistically significant in comparison to free romidepsin.

**Figure 7.**
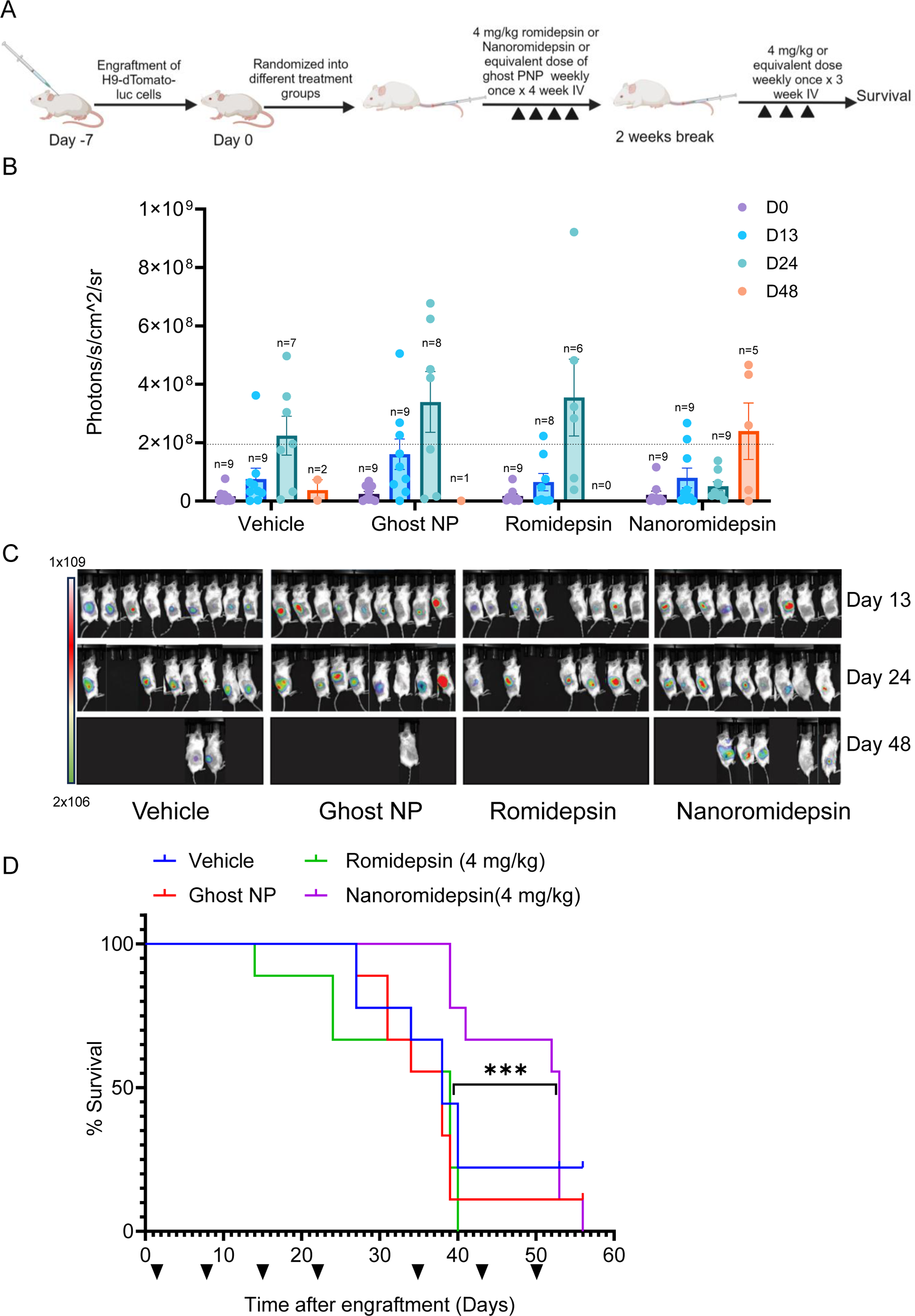
Dosing schedule change of Romidepsin encapsulated nanoparticle showed superior activity and survival rate compared to free romidepsin in CTCL xenograft bearing NSG mice. (A) Diagram representing the inoculation and dosing schedule of free romidepsin and Nanoromidepsin in TCL xenograft bearing mice; (B) Region-of-interest analysis of BLIs (readout for tumor growth) from different treatment groups at various time points during the course of treatment and plotted as bar graph; (C) Whole-body bioluminescence images of H9-dTomato-luc xenograft–bearing mice taken at the indicated day; (D) Survival curves for romidepsin–treated, Nanoromidepsin-treated, ghost nanoparticle and control mice (n = 9 per group). The arrows indicate treatment days. Statistical significance was determined by using log rank test where *, p<0.05, **< p<0.01, ***, p<0.0001.

## Discussion

The dwindling options to treat patients with relapsed or refractory PTCL have created an urgent need to change the paradigm in how we think about and develop new drugs for challenging orphan diseases. In the U.S., pralatrexate and the HDACi belinostat are the only drugs still approved for patients with R/R PTCL, albeit they have only a dangling accelerated approval.

Brentuximab vedotin (Bv), an antibody drug conjugate which targets CD30, has been approved for the treatment of R/R anaplastic large cell lymphoma (ALCL), and in combination with cyclophosphamide, doxorubicin and prednisone (Bv-CHP) for the front-line treatment of CD30-positive PTCL(*33*). Most physicians who treat patients with R/R PTCL agree that romidepsin is among one of the more important drugs they prescribe to manage these diseases. Loss of the romidepsin indication in R/R PTCL has put physicians and patients in a challenging position. Couple this with the reality that there are not many new drugs on the horizon, it becomes obvious that new strategies to improve the agents at our disposal, or to create new ones based on promising combinatorial datasets, represent one relatively derisked approach to advance care.

Romidepsin in combination with other epigenetically targeted drugs like the DNMT3 inhibitor 5-azacytidine appears to produce the best overall response rate (ORR) and progression free survival (PFS) data of any ‘drug’ or drug combination to date in this population (*34, 35*). Nonetheless, these modest clinical and preclinical experiences suggest that combinations with an HDACi, with romidepsin being among the most potent in the class, may represent one relatively straight-forward path to create new treatment platforms for this challenging population. How these combinations get selected may have an impact on the clinical experiences, as those doublets with robust preclinical datasets and a demonstration of drug : drug synergy appear to have generally more favorable outcomes (*36, 37*). While romidepsin has been shown to be consistently more potent than other HDACi in the laboratory, in the clinic it produces an ORR of 25%, with a PFS of about 3 to 4 months, and a median duration of response of more than a year (*5, 38*). These discordant findings between pre-clinical and clinical experiences remain to be explained. Pharmacokinetic analysis suggests that romidepsin is highly protein-bound (92%-94%), with α1-acid-glycoprotein (AAG) being the principal binding protein. Also, an *in vitro* study showed that romidepsin accumulates in human hepatocytes via an unknown active uptake process (ISTODAX package insert), a finding we validated in the murine model system. After administration of 4 mg/kg free romidepsin and Nanoromidepsin weekly for three weeks in mice, LC-MS based quantification of romidepsin in liver demonstrated greater accumulation of free romidepsin, while mice treated with Nanoromidepsin exhibited substantially less accumulation of romidepsin in the liver (Figure 5H). One possible explanation for why all HDAC inhibitors produce similar clinical data despite vast differences in potency may relate to the sub-optimized pharmacologic features of the drug, and the fact that the short half-life (estimated to be about 3.8 hours in humans), coupled to a high degree of protein binding and a smaller volume of distribution (Vd) may not maximize romidepsin’s effects on transcriptional activation, the primary mechanism of action for the drug. Strategies that optimize the on-target effect, transcription, offer some prospect of improving the effect of any epigenetic based drug. Here, we exploited the unique physicochemical properties of a tailored PNP, including optimal size and surface properties, enhanced volume of distribution (Vd), and augmented tumor bioavailability, in an effort to improve on the fundamental mechanism of action of the epigenetic effect. All of these factors are essential to overcome those factors that might compromise drug activity. The convergence of the drugs’ mechanism of action and the features of a PNP offer the prospect of optimizing the anti-tumor effects of the HDACi.

Polymer-based nanoparticles are receiving increasing attention because of their unique properties that dramatically improve many of the liabilities associated with sub-optimized drugs (*39*). Amphiphilic block co-polymers offer many advantages over traditional nanoliposomal based technologies, largely because they can accommodate a broader range of drugs with variable solubility features, that is, both lipophilic and hydrophilic molecules. Polymer nanoparticles typically have a size of less than 100 nm, about the size of a virus, which aids in improving the circulation time of the drug and the volume of distribution, allowing for a bioconcentration of drug in tissue, particularly tumor tissue. The bioluminescent *in vivo* assay which deployed a PNP containing both romidepsin and DIO clearly establishes a predilection for the PNP to bioaccumulate in the tumor microenvironment. Although there is some debate about the precise mechanisms by which polymeric nanoparticles accumulate in the tumor microenvironment, porous and leaky vasculature have been advanced as one of the explanations (*40*). Irrespective of the underlying mechanics, we have clearly shown a penchant for drug to bioaccumulate in tumor. Generally, a variety of polymers have been deployed for the synthesis of PNPs, including polylactides (PLA) (*41, 42*), poly (lactide co-glycolides) (PLGA) (*43, 44*), and poly(D,L-Lactic Acid) (*45*). These polymers are considered biocompatible, biodegradable and non-toxic, which enhances their elimination, improves their tolerability, and reduces their immunogenicity (*46*). Another important component of the PNP is the inclusion of the PEG chain which has been shown to reduce the elimination of the particles via the host immune system and thereby maximizes the circulation time (*47*). An attractive feature of this platform is that hydrophobic drugs can be readily incorporated and even conjugated to the polymer (*48–50*). Surface conjugation of cell specific targeting ligands to PEG can also be performed (*51–53*).

Based upon the clear liabilities associated with the present use of romidepsin, and at least the theoretical advantages of leveraging a nanochemistry platform, we initiated a program to synthesize a polymer nanoparticle of romidepsin to overcome some of these well-established liabilities of the drug, while intently focused on trying to improve romidepsin’s fundamental epigenetic mechanisms of action to alter gene expression (*54, 55*). PNPs of romidepsin were synthesized using FDA approved GRAS components. We systematically investigated the parameter effects on the physicochemical properties of the nanoparticles, in particular the encapsulation efficiency of the drug. We optimized a precise and scalable method of nanoparticle synthesis by using widely accessible GRAS materials, a crucial step for translational therapeutic development (*56*).

In general, the physicochemical data suggested that the nanoprecipitation technique optimized the desired parameters of the nanoparticle. DLS data showed the unimodal distribution of the particles, with a PDI of 0.145 with z-average of 46.25 nm in size. We designed these particles to be approximately 50 nm, which has been suggested to be a size feature that facilitates bioconcentration in the tumor microenvironment (*57*). Particles in this size range are thought to extravasate into neovascularized tumor tissue. By optimizing the pharmacologic features of the romidepsin PNP, we demonstrated superior potency *in vitro* compared to free romidepsin in different subtypes of T-cell lymphoma and LGL leukemia cell lines. In addition, Nanoromidepsin induced apoptosis as evidenced by the increase in cleaved PARP similar to free romidepsin, which is aligned with prior evidence.

In patients with T-cell lymphomas, administration of romidepsin at a dose of 14 mg/m2 intravenously over 4-hours on days 1,8, and 15 of a 28-day cycle results in geometric mean values of maximum plasma concentration (C_max_) and area under the plasma concentration-time curve (AUC0-∞) of 377 ng/mL and 1549 ng*hr/mL, respectively. Romidepsin administered as a slow IV bolus dose to rats at 0.33 and 0.67 mg/kg (Report 501650; Istodax package insert) achieved a mean AUC∞ of 10.3 and 18.1 ng.hr/mL, respectively, after a single dose. (Istodax package insert). Recognizing all the cross-species differences, these data suggest that Nanoromidepsin in these murine models approximated or dramatically exceeded those PK parameters established in humans. Following a single intravenous dose of romidepsin and Nanoromidepsin, the C_max_ was 21.31 and 231 ng/ml respectively. Another major difference was seen in the AUC, which was 231 and 2532 ng/mL for free and Nanoromidepsin respectively. Nanoromidepsin also exhibited a 1.5-fold increase in half-life (t1/2) compared to free romidepsin, indicating prolonged availability in plasma. These PK features of Nanoromidepsin were confirmed in the biodistribution study where Nanoromidepsin was shown to preferentially bioaccumulate in the tumor over other vital organs. This is noteworthy as some conventional polymeric nanoparticles have been shown to accumulate in organs like the spleen, liver, and kidneys, potentially limiting their therapeutic potential (*47*). These findings are concordant with previous studies indicating that a PNP tailored for the API can improve bioavailability and solubility issues, thereby optimizing their mechanism of action (*58, 59*), a factor that may be especially important for drugs targeting the epigenome.

The improvement in the PK parameters supporting improved drug exposure of course raises concerns about incrementally worse tolerability. As shown in a series of comprehensive single and multiple dose toxicity assays, Nanoromidepsin was found to be substantially safer than free romidepsin, even at the highest doses studied. These data have established a sound basis to identify the MTD, optimal route of administration, and ideal dosing schedule prior to the efficacy studies. The *in vivo* toxicity assays affirmed that Nanoromidepsin was safer compared to free romidepsin, exhibiting less accumulation in the liver as observed in the biodistribution studies, and as supported by the histopathology and quantitative LC-MS-based quantification of drugs in organs. In a TCL xenograft model, Nanoromidepsin exhibited an LD50 value of 8 mg/kg, while free romidepsin exhibited an LD50 value of 5 mg/kg. Clinically, the most commonly reported hematologic adverse effects of romidepsin include thrombocytopenia, anemia and neutropenia, while the major non-hematologic toxicities include asthenia, infection, and GI disturbance (*60*). The direct comparison of body weight loss and clinical toxicity scores in mice confirmed the superior safety profile of Nanoromidepsin at every dose and schedule studied.

Across all efficacy studies performed, Nanoromidepsin produced substantially superior growth delay, and an overall survival advantage in contrast to free romidepsin. Generally, multiple cycles of drug administration are uncommonly explored in *in vivo* murine studies, despite it being the standard of care in human patients. This difference makes it difficult to see overall survival advantages in murine models. An overall survival advantage is based on the depth of a complete remission (CR), which in normal clinical practice is usually achieved with multiple cycles of combination therapy. The improved tolerability and efficacy of Nanoromidepsin would suggest that combinations of drugs with Nanoromidepsin will further deepen the CR, likely translating into improved outcomes for patients with PTCL.

In summary, we have pioneered the development of a unique romidepsin polymer nanoparticle, which exhibits superior pharmacokinetic features, tolerability, and efficacy compared to the historically approved drug. The improved effects on transcription may explain the improvement in efficacy and is supported by the biodistribution data confirming marked bioaccumulation in the tumor microenvironment. This study represents the first to interrogate the merits of a polymer nanoparticle platform on the pharmacology and activity of an epigenetically targeted drug in these diseases. Future studies will address the mechanisms that account for the bioaccumulation of the romidepsin PNP in the tumor microenvironment, as well as the differences in gene expression and how this might explain the potent efficacy advantage for Nanoromidepsin. We believe the platform has potentially created an opportunity to reconfigure the traditional treatment paradigms for patients with peripheral T-cell lymphoma, as we now poise this drug for future clinical studies.

## Acknowledgements

This manuscript is dedicated to the memory of Dr. Mark Kester, a colleague, friend, and pioneer in the field of nanochemistry. We acknowledge Jeremy Gatesman, Joshua Tennant and Susan Walker for their contribution. We acknowledge Molecular Immunologic & Translational Sciences (MITS) Core for animal work.

OAO is grateful for the American Cancer Society Research Professorship. OAO is funded in part by Grant 1R01FD006814-01. The LC-MS work is supported by P30 CA044579 (TEF). The authors are indebted to the Scarlet Feather Foundation and the IVY Foundation for their continued support. The authors thank University of Virginia LGL Leukemia Registry personnel for their support of this study. The Registry was supported by the Bess Family Charitable fund and a gift from a generous anonymous donor (TPL). The data for this (manuscript or presentation) were generated in the University of Virginia Flow Cytometry Core Facility (RRid:SCR_017829) and is partially supported by the NCI Grant (P30-CA044579). Cell sorting was performed on the Cytek Aurora CS funded through the NIH S10 instrumentation grant (1S10OD034355-01).

