## Supplemental Figures for "Preclinical Development of a Romidepsin Nanoparticle Demonstrates Superior Tolerability and Efficacy in Models of Human T-Cell Lymphoma and Large Granular Lymphocyte Leukemia"

### Slide 1
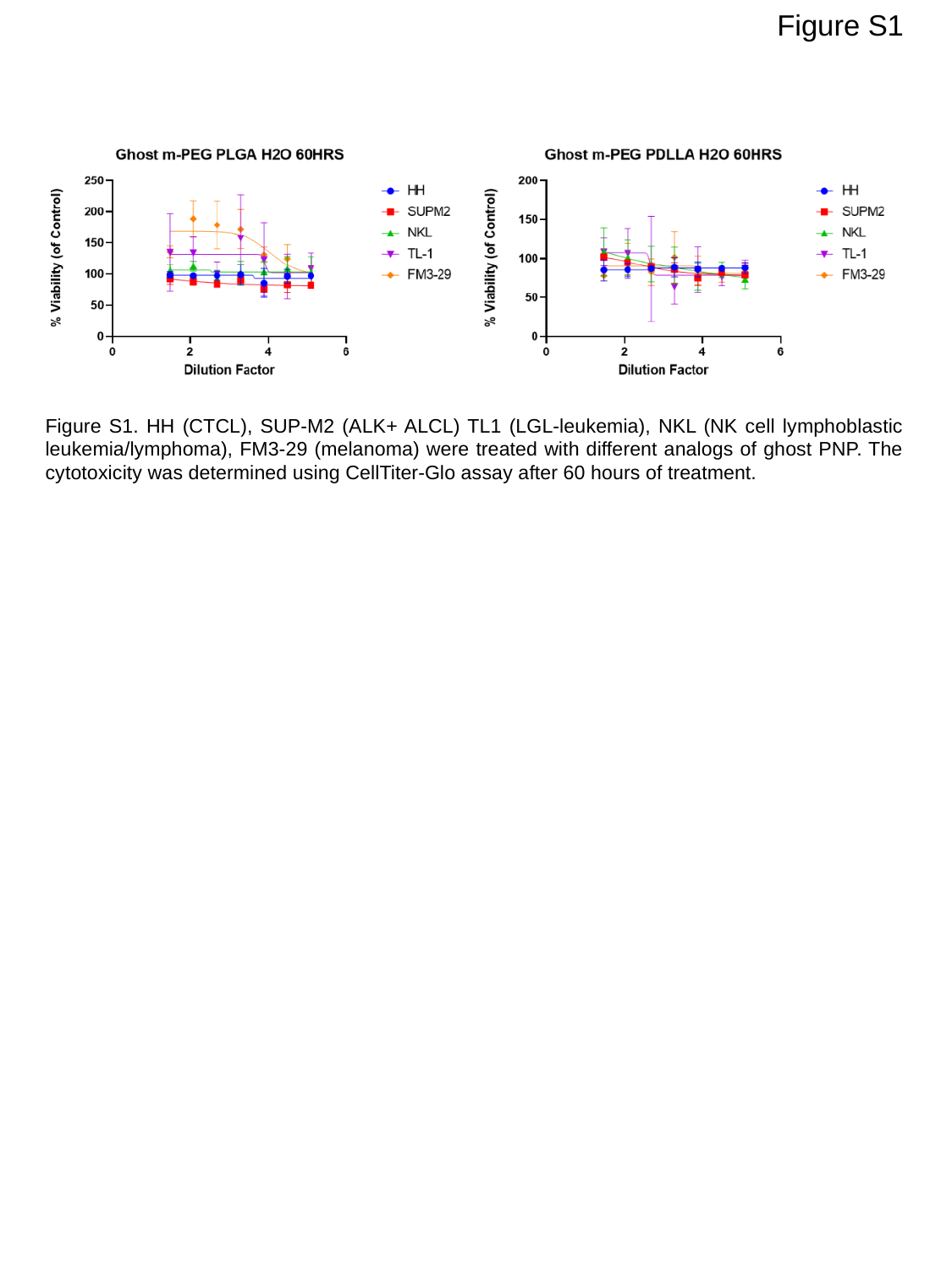

Figure S1
Figure S1. HH (CTCL), SUP-M2 (ALK+ ALCL) TL1 (LGL-leukemia), NKL (NK cell lymphoblastic leukemia/lymphoma), FM3-29 (melanoma) were treated with different analogs of ghost PNP. The cytotoxicity was determined using CellTiter-Glo assay after 60 hours of treatment.

### Slide 2
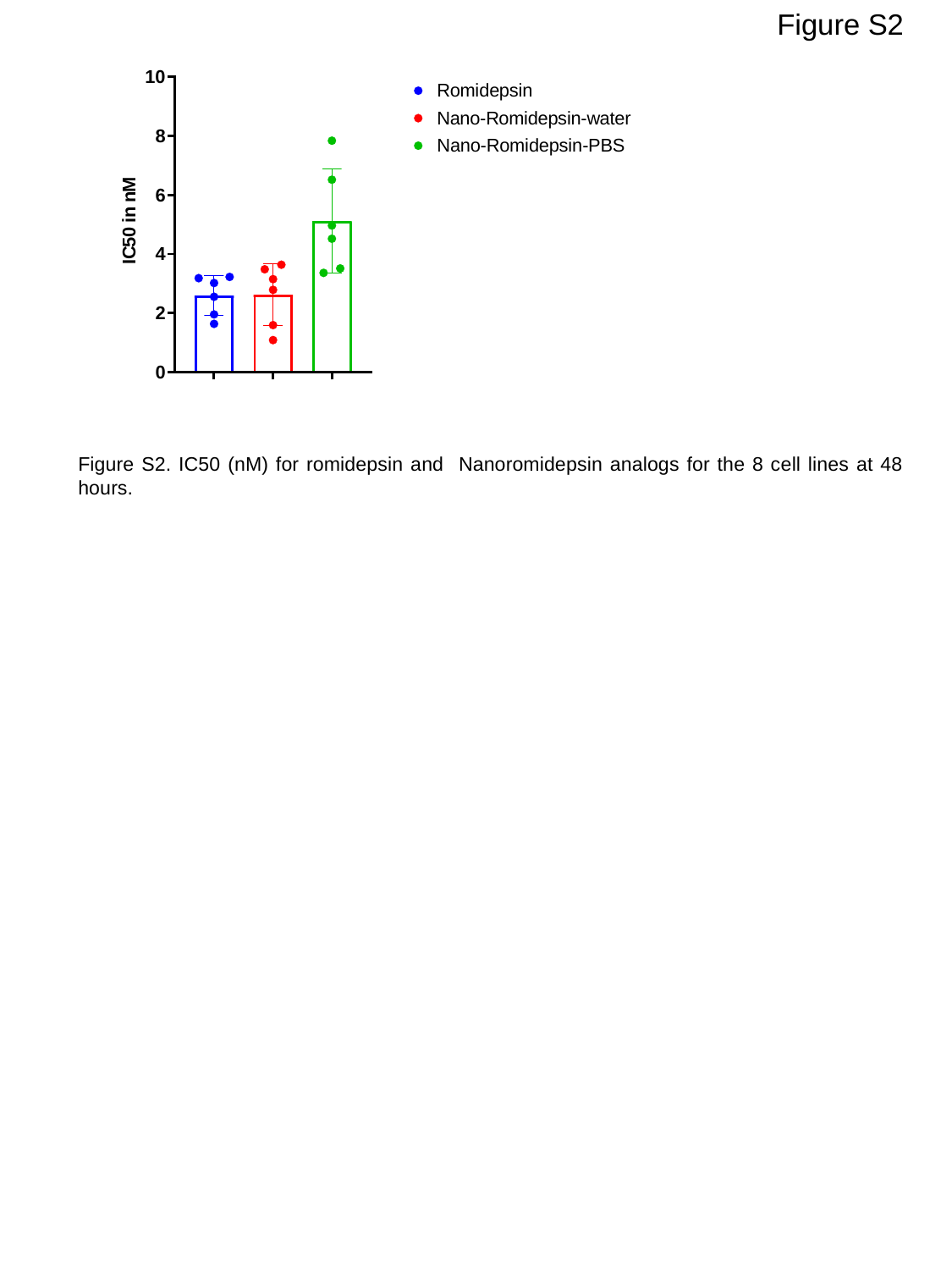

Figure S2
Figure S2. IC50 (nM) for romidepsin and Nanoromidepsin analogs for the 8 cell lines at 48 hours.

### Slide 3
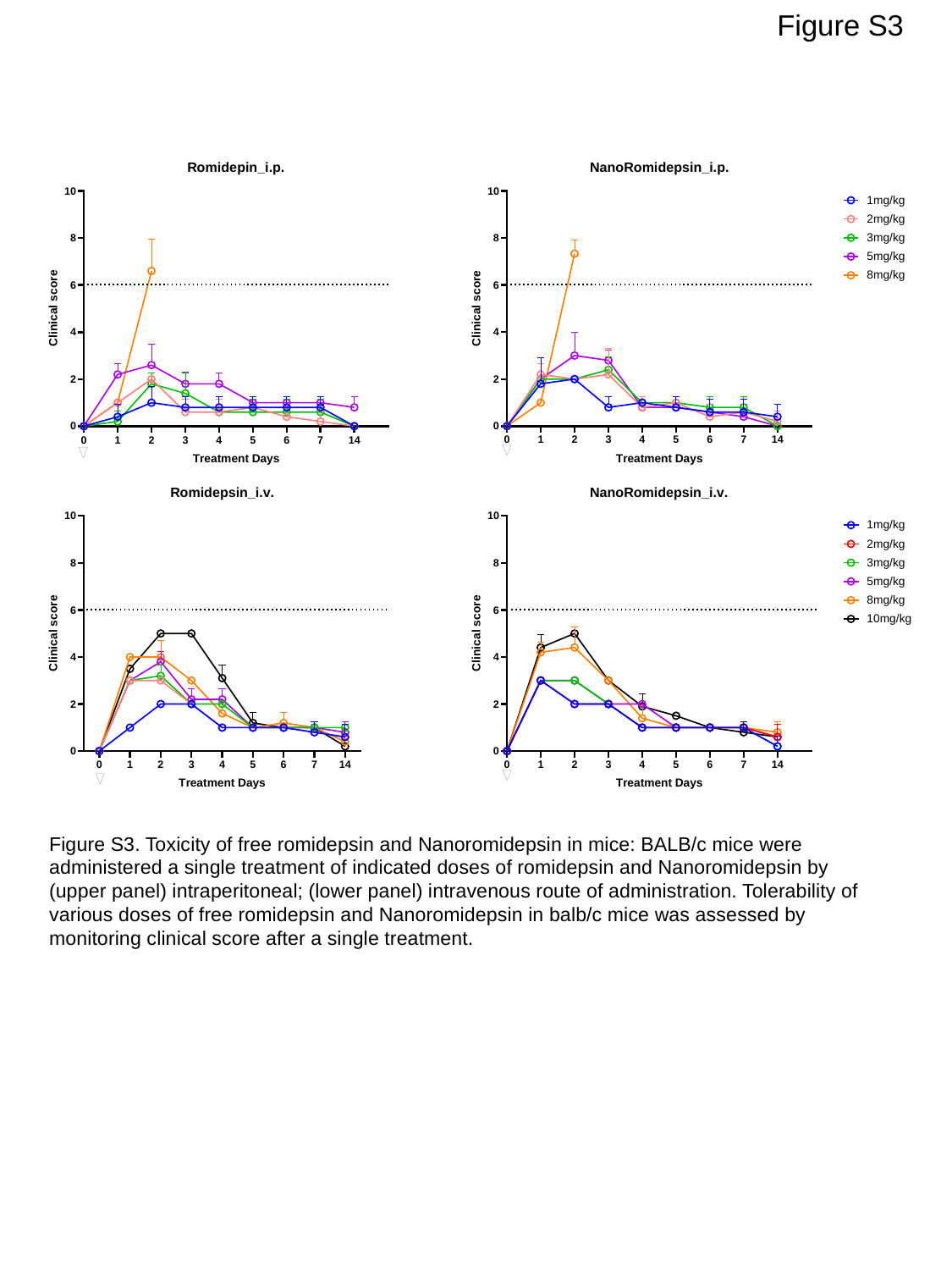

Figure S3
Figure S3. Toxicity of free romidepsin and Nanoromidepsin in mice: BALB/c mice were administered a single treatment of indicated doses of romidepsin and Nanoromidepsin by (upper panel) intraperitoneal; (lower panel) intravenous route of administration. Tolerability of various doses of free romidepsin and Nanoromidepsin in balb/c mice was assessed by monitoring clinical score after a single treatment.

### Slide 4
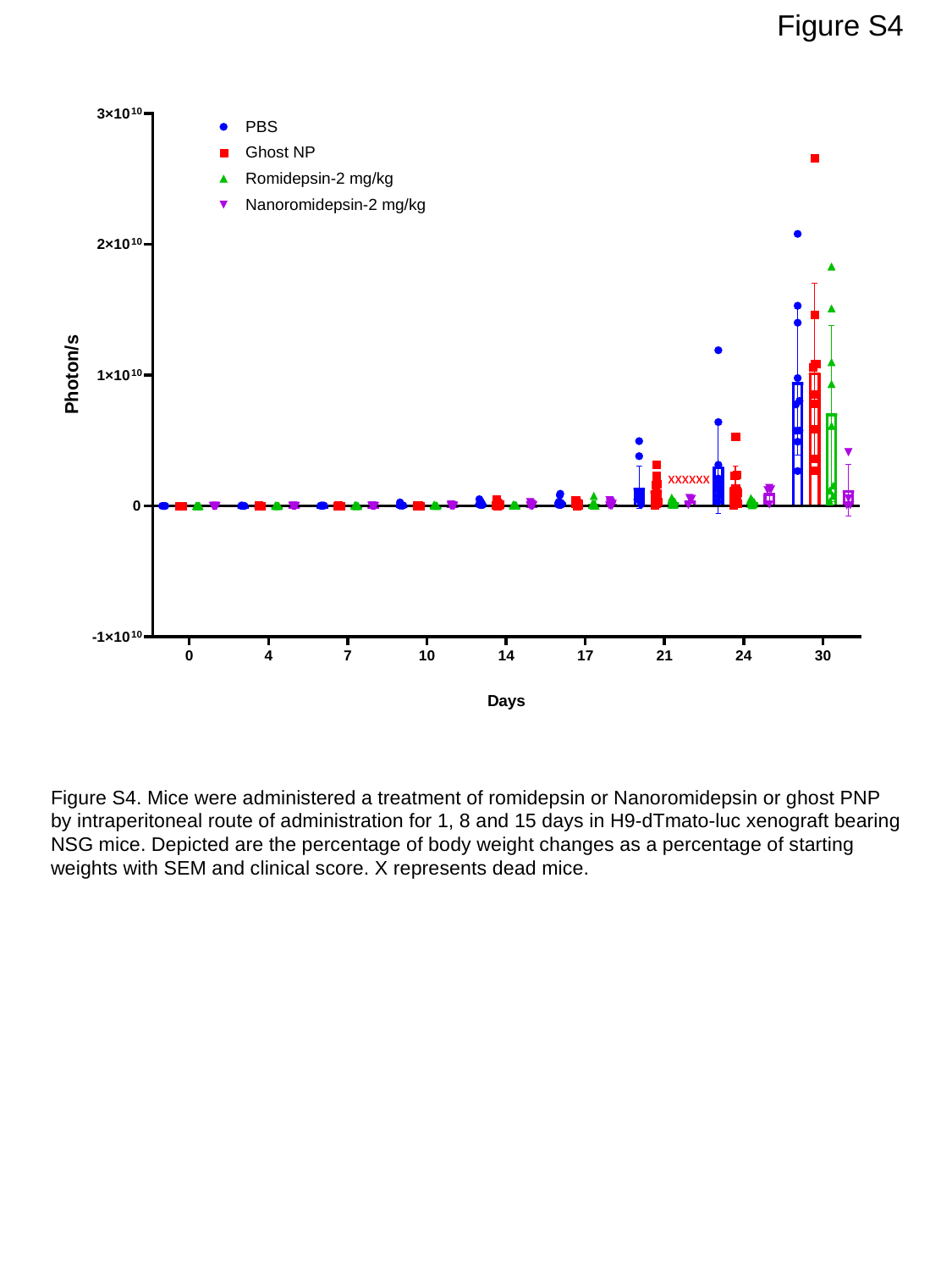

Figure S4
xxxxxx
Figure S4. Mice were administered a treatment of romidepsin or Nanoromidepsin or ghost PNP by intraperitoneal route of administration for 1, 8 and 15 days in H9-dTmato-luc xenograft bearing NSG mice. Depicted are the percentage of body weight changes as a percentage of starting weights with SEM and clinical score. X represents dead mice.

### Slide 5
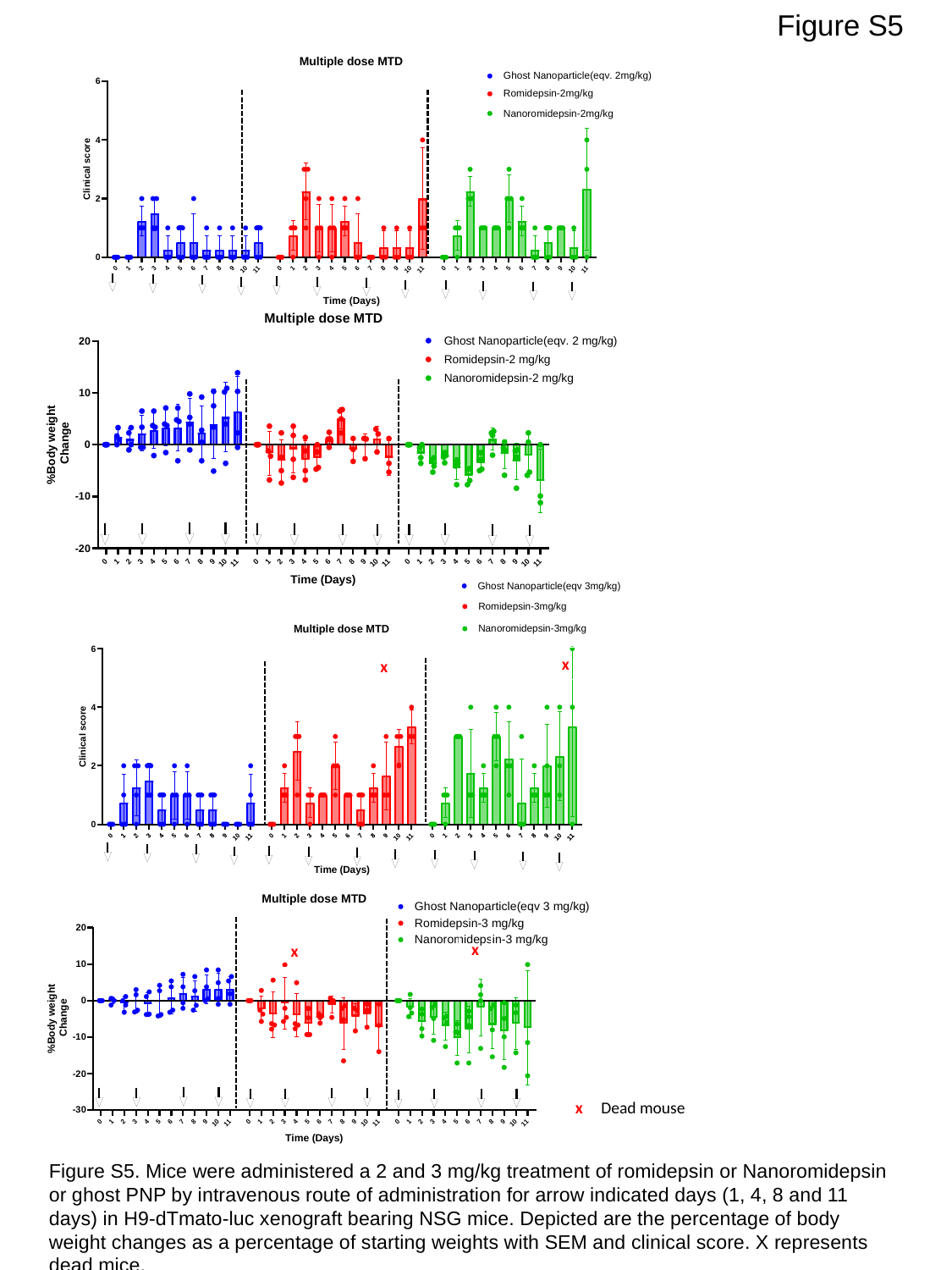

Figure S5
x
x
x
x
x
Dead mouse
Figure S5. Mice were administered a 2 and 3 mg/kg treatment of romidepsin or Nanoromidepsin or ghost PNP by intravenous route of administration for arrow indicated days (1, 4, 8 and 11 days) in H9-dTmato-luc xenograft bearing NSG mice. Depicted are the percentage of body weight changes as a percentage of starting weights with SEM and clinical score. X represents dead mice.

### Slide 6
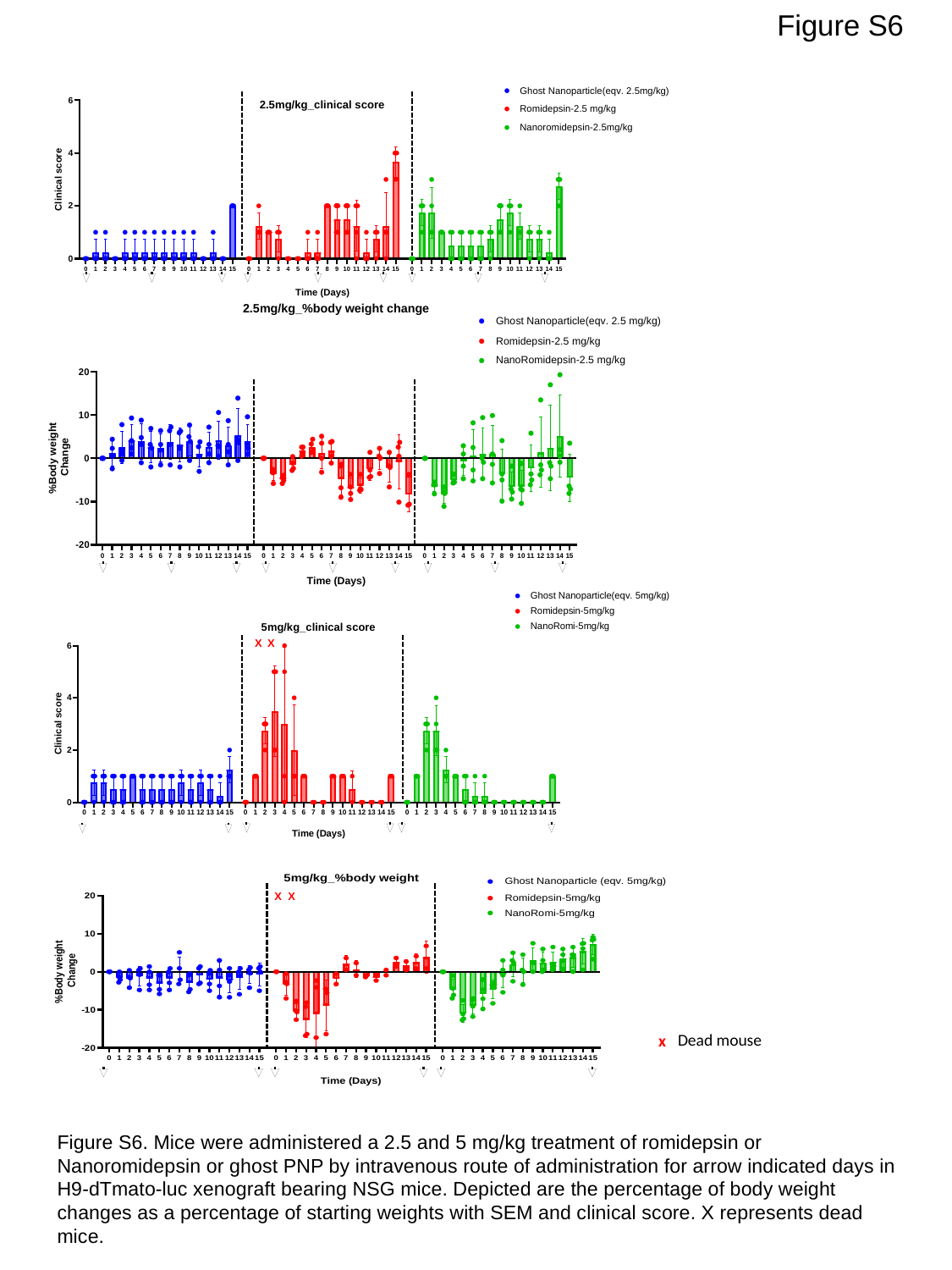

Figure S6
X
X
X
X
Dead mouse
x
Figure S6. Mice were administered a 2.5 and 5 mg/kg treatment of romidepsin or Nanoromidepsin or ghost PNP by intravenous route of administration for arrow indicated days in H9-dTmato-luc xenograft bearing NSG mice. Depicted are the percentage of body weight changes as a percentage of starting weights with SEM and clinical score. X represents dead mice.
